# Unveiling the Evolution of Antimicrobial Peptides in Gut Microbes via Foundation Model-Powered Framework

**DOI:** 10.1101/2025.01.13.632881

**Authors:** Wenhui Li, Baicheng Huang, Menghao Guo, Zihan Zeng, Tao Cai, Linqing Feng, Xinpeng Zhang, Ling Guo, Xianyue Jiang, Yanbin Yin, Ercheng Wang, Xingxu Huang, Jinfang Zheng

**Affiliations:** Research Center for Life Sciences Computing, Zhejiang Lab, Hangzhou 311121, Zhejiang, China; School of Life Sciences and Technology, Tongji University, Shanghai 200092, China; Nebraska Food for Health Center, Department of Food Science and Technology, University of Nebraska, Lincoln, NE 68588, USA; Zhejiang Provincial Key Laboratory of Pancreatic Disease, The First Affiliated Hospital, and Institute of Translational Medicine, Zhejiang University School of Medicine, Hangzhou 311121, China; School of Life Science and Technology, ShanghaiTech University, Shanghai 200092, China

## Abstract

Antimicrobial resistance poses a growing threat to public health, emphasizing the urgent need for novel therapeutic strategies. Antimicrobial peptides (AMPs), short peptide sequences with diverse mechanisms of action, offer a promising alternative due to their broad-spectrum activity against pathogens. Recent advances in protein language models (PLMs) have revolutionized protein structure prediction and functional annotation, highlighting their potential for AMP discovery and therapeutic development. In this context, we present AMP-SEMiner (Antimicrobial Peptide Structural Evolution Miner), an AI-driven framework designed to identify AMPs from metagenome-assembled genomes (MAGs). By integrating PLMs, structural clustering and evolutionary analysis into the framework, AMP-SEMiner can identify AMPs encoded by small open reading frames (smORFs) and encrypted peptides (EPs), significantly expanding the discovery space. Using this approach, we identified 1,670,600 AMP candidates from diverse habitats. Experimental validation of 29 candidates revealed antimicrobial activity in 18, with 13 surpassing antibiotics in effectiveness. Further analysis of AMPs from human gut microbiomes demonstrated both conserved and adaptive evolutionary strategies, ensuring their functional efficacy in the dynamic gut environment. These findings position AMP-SEMiner as a powerful tool for the discovery and characterization of novel AMPs, with significant potential to drive the development of new antimicrobial therapies.

## Introduction

Antibiotic-resistant infections are becoming increasingly challenging to treat with conventional therapies, highlighting the urgent need for novel antibiotic^1^. In this context, antimicrobial peptides (AMPs), short chains of 5 to 100 amino acids found in all organisms, are emerging as a promising alternative due to their unique mechanisms against microbes^2^. The antimicrobial mechanisms of AMPs include disrupting microbial cell walls^3^, cell membranes^4^, or targeting intracellular components^5^. These mechanisms allow AMPs to address a broad spectrum of microbes, including bacteria, fungi^6^ and even cancer cells^7^, which distinguishes them from traditional antibiotics. Naturally, AMPs are produced through processes such as proteolytic cleavage from encrypted peptides (EPs)^8^, non-ribosomal synthesis^9^, or direct genome encoding^10^.

Recent studies have identified numerous novel AMPs from genomic and metagenomic data, complementing traditional wet-lab methods. These AMPs, sourced from animal^11–13^, human^14, 15^, and soil microbiomes^16^, as well as de-extinct human genomes^17^, show promise as therapeutics. Naturally occurring AMPs from the human genome and symbiotic microbes are expected to have low toxicity and mild antimicrobial activity, helping maintain microbiota balance for long-term health^18, 19^. Advances in metagenomic sequencing have expanded the search space, increasing the potential for discovering new AMPs with unique therapeutic applications while preserving microbiota balance.

Computational approaches for identifying AMPs typically employ two main strategies: (1) Protein Classification, which frames AMP detection as a protein sequence classification task. While effective, these methods^20–24^ mainly identified small open reading frames (smORFs) and may miss AMPs that exist as fragments within larger proteins, such as Cathelicidin^25^. (2) Cleavage Site Prediction, exemplified by tools like panCleave^17^, focuses on identifying protease cleavage sites to generate protein fragments as AMP candidates. However, this method is mainly optimized for human proteins and does not directly identify AMPs in microbial proteins or fragments. Recently, the development of self-supervised protein language models (PLMs) such as ESM2^26^ and ProtT5^27^ marks a significant advancement in protein science. Models trained on large datasets excel in applications such as protein function annotation^28, 29^ and 3D structure prediction^26^. Given their effectiveness and the similarities to AMP discovery, PLMs are expected to enhance AMP mining due to the following advantages: enhanced representation capacity and incorporation of structural insights.

In this study, we developed AMP-SEMiner (**A**nti**m**icrobial **P**eptide **S**tructural **E**volution **Miner**), a robust and comprehensive AI framework for identifying AMP from metagenome-assembled genomes (MAGs). This framework integrates PLM, structural clustering, evolutionary analysis, experimental validation, and molecular dynamics (MD) simulations to explore AMP mechanisms of action. Through methodological innovations, AMP-SEMiner enables the identification of AMPs encoded by both smORFs and EPs, expanding the discovery space. Using this framework, we identified 1,670,600 predicted AMPs from diverse habitats. Out of 29 AMP candidates experimentally tested, 18 exhibited antimicrobial activity, with 13 outperforming antibiotics. Further analysis of AMPs from four human gut microbiomes showed that AMP evolution combines conserved and adaptive strategies, allowing functional efficacy while adapting to the dynamic gut environment. These findings highlight AMP-SEMiner as a powerful tool for discovering novel AMPs and advancing antimicrobial therapy development.

## Results

### 1 A comprehensive AI framework for AMP mining

We developed AMP-SEMiner, a unified AI framework for the simultaneous identification of AMPs as both smORFs and EP antibiotics. The framework comprises five key components (**Fig. 1**):

**Fig. 1.**
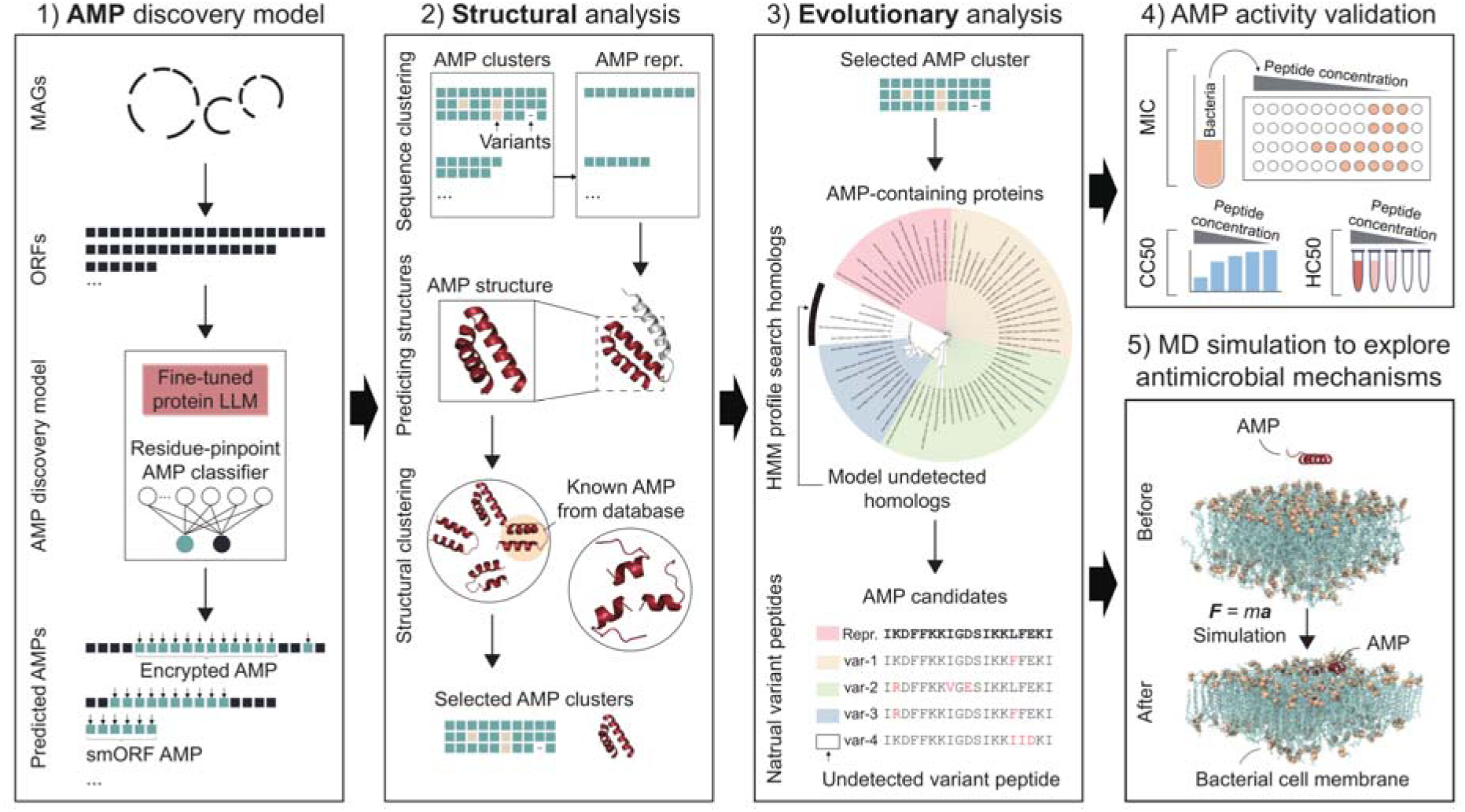
Schematic overview of the AMP-SEMiner framework. This diagram outlines the five key components of the AMP-SEMiner framework: (1) **AMP discovery model**: Utilizes a protein foundation model for residue-level prediction to identify potential AMPs. (2) **Structural analysis**: Applies sequence and structural clustering to categorize the identified AMPs into distinct groups. (3) **Evolutionary analysis**: Identifies homologs of AMP-containing proteins and their natural variants as potential AMP candidates. (4) **Experimental validation**: Assesses the antimicrobial activity of candidates, along with their cytotoxicity, hemolytic properties and propidium iodide staining. (5) **Molecular dynamics (MD) simulation**: Investigates the mechanisms of action of active AMPs through computational modeling.

#### AMP discovery model

This model takes proteins from MAGs, integrating a fine-tuned PLM with a residue-level AMP classifier. It predicts the likelihood of each residue being AMP or non-AMP. Continuous sequences of AMP residues are identified as potential AMPs (>= 5aa). This process enables the discovery of AMP from both smORFs and EPs.

#### Structural analysis

Predicted AMPs undergo sequence clustering and 3D structure prediction via AlphaFold2^30^. Structural clustering with FoldSeek^31^, using both predicted and known AMP structures (e.g., from APD3^32^), refines the analysis. This step facilitates the selection of AMP clusters based on sequence/structure cluster-size and physicochemical properties, linking sequence, structure, and function.

#### Evolutionary analysis

Evolutionary analysis of selected AMP clusters involves constructing Hidden Markov Model (HMM) profiles and searching metagenomic ORFs with HMMER^33^. This approach uncovers homologs and natural variants, identifying entire AMP families as potential candidates.

#### Experimental validation of AMP

The antimicrobial activity of the AMPs was evaluated by determining their minimum inhibitory concentration (MIC) against both Gram-positive and Gram-negative bacteria. To assess their safety profile, the 50% cytotoxic concentration (CC50) and 50% hemolytic concentration (HC50) were measured in mammalian cells. Propidium iodide (PI) staining is used to investigate the mechanism of action.

#### Computational validation of AMP-membrane interactions

All-atom molecular dynamics simulations were utilized to explore the dynamics of AMPs as they transitioned from positions distant from the membrane surface to engaging in interactions with the membrane. Furthermore, energy decomposition analysis was performed to pinpoint residues with significant energetic contributions and to identify the dominant energetic components driving these interactions.

### 2 Advancing AMP-SEMiner discovery model through dataset construction, model optimization and comprehensive benchmarking

#### 2.1 Construction of residue-level AMP datasets to overcome length bias

The distinct patterns exhibited by known AMPs^34, 35^ and background proteins pose a length distribution challenge **(Fig. 2a**). To address this, we propose a data preprocessing strategy that maps AMP to the source proteins, thereby constructing the AMP-positive dataset with all sequences labeled at the residue level (**Fig. 2b and Methods**). Background proteins were designated as the negative data. This approach yielded 18,992 AMP-containing proteins and 2,106,517 negative proteins, with similar sequence length distributions to mitigate bias in the dataset (**Fig. 2c**).

**Fig. 2.**
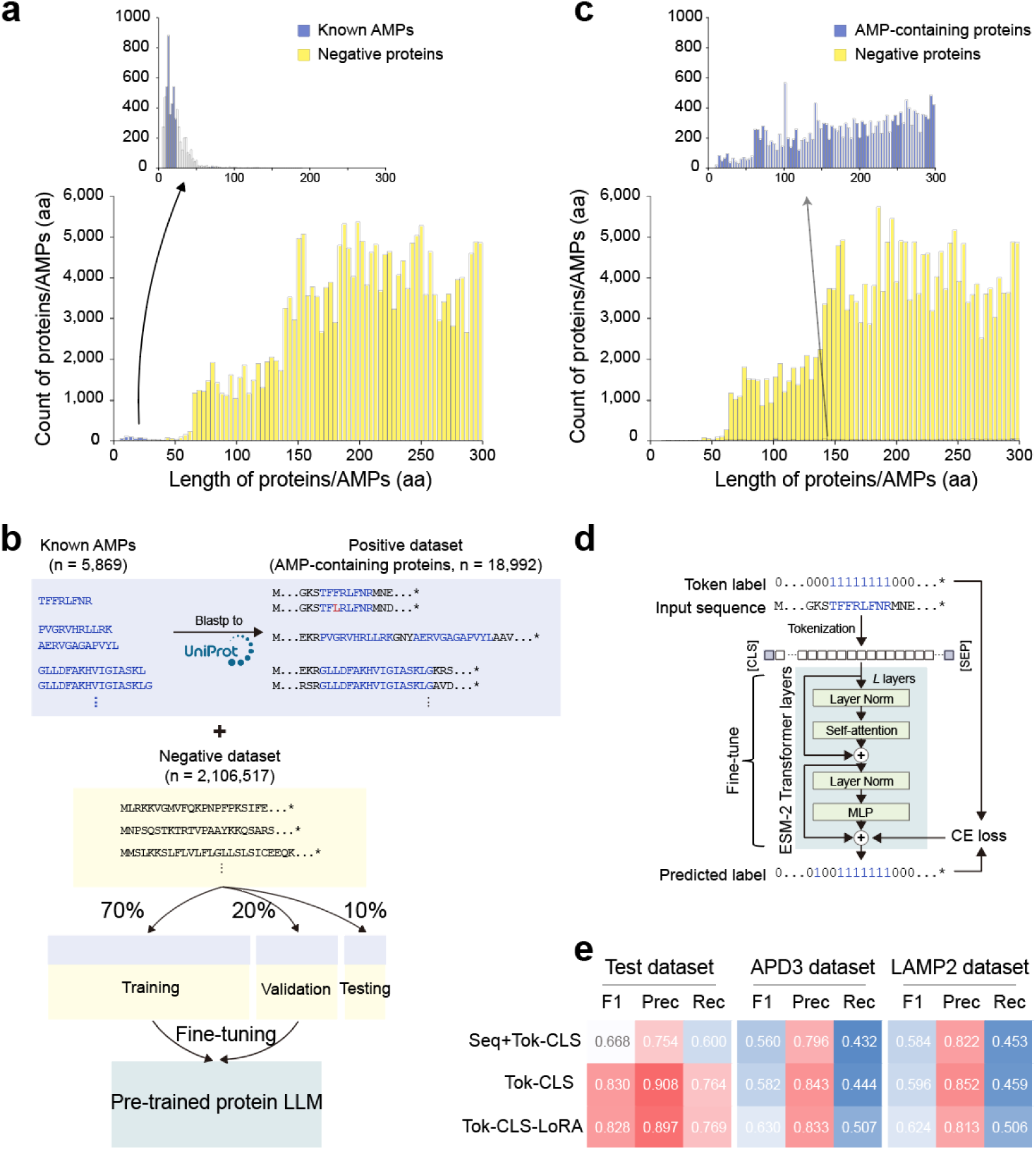
Overview of the AMP-SEMiner discovery model. **(a)** Length distributions between known AMPs and background proteins, highlighting the length bias in previous methods. **(b)** Workflow for processing AMP sequence data and constructing the datasets. **(c)** Distribution of sequence lengths in our dataset, where the positive and negative data show similar length distributions, effectively mitigating the length bias in the model. **(d)** Overview of the model training strategy for residue-level prediction of potential AMPs. **(e)** Performance evaluation on the testing dataset and two independent datasets (APD3 and LAMP2), assessed at the residue level, demonstrating the high accuracy and robustness of AMP-SEMiner discovery model. (1) Seq+Tok-CLS: a two-step approach combining a fine-tuned ESM-2 for sequence classification and a fine-tuned ESM-2 for token classification. (2) Tok-CLS: an end-to-end fine-tuned ESM-2 for token classification (‘Tok-CLS’), and (3) Tok-CLS-LoRA: an end-to-end fine-tuned ESM-2 with Low-Rank Adaptation (LoRA) for token classification.

#### 2.2 Fine-tuning ESM-2 with optimized parameters for AMP discovery at residue level

To train models for residue-level identification of AMPs, we conducted extensive experiments to optimize model parameters (**Supplementary Fig. 1** and **Supplementary Tables 1-6)**. Ultimately, we adopted an end-to-end token classification approach, fine-tuning the protein foundation model ESM-2 using a cross-entropy (CE) loss function. Each residue in AMP-containing and negative proteins was labeled as either 1 or 0, representing AMP-positive and AMP-negative tokens, respectively (**Fig. 2d**). The tokenized data were feed into the ESM-2, followed by a two-layer linear classifier implemented as a multilayer perceptron (MLP). Both the ESM-2 Transformer layers and the classifier were jointly trained to minimize token-level CE loss (**see Methods**).

#### 2.3 Benchmarking of AMP discovery approaches at protein and residue level

To evaluate the performance of our AMP discovery approach, we conducted benchmarking against several published AMP identification algorithms at both the protein and residue level. In addition to our testing dataset, we created two independent datasets based on the APD3 and LAMP2^36^. Our benchmarking includes over half of the tools published since 2017 (**Supplementary Table 7**).

For protein level, the performance of various tools is summarized in **Table 1**. Pre-trained protein language model-based methods, such as Ma et al. (2022)^21^ and AMP-BERT, generally outperformed traditional machine learning approaches but displayed a trade-off between precision and recall. Ma et al. (2022) achieved high precision but low recall due to strict filtering that excluded some true AMPs. In contrast, AMP-BERT had high recall but low precision, reflecting a higher false positive rate and revealing the limitations of relying solely on PLMs for sequence classification. Models combining sequence and structural data, like PGAT-ABPp and deepAMPNet, underperformed relative to other methods. However, our model demonstrated superior performance in identifying AMP-containing proteins across the testing and independent datasets, primarily due to a data preprocessing strategy that broadened the AMP discovery scope without focusing solely on smORF detection.

**Table 1.** Performance of AMP-containing protein identification by various approaches on independent test datasets.

| Approach | Published Year | Model Type | Input Data type | Test dataset (+1880/-208546) |  |  | APD3 dataset (+5941) | LAMP2 dataset (+5498) |
| --- | --- | --- | --- | --- | --- | --- | --- | --- |
|  |  |  |  | F1 | Precision | Recall | Recall | Recall |
| Macrel<sup>20</sup> | 2020 | SVM, KNN, LR, DT, RF | Seq. | 0.044 | 0.086 | 0.030 | 0.041 | 0.047 |
| AmpGram<sup>37</sup> | 2020 | RF | Seq. | 0.035 | 0.018 | 0.545 | **0.626** | 0.665 |
| ampir<sup>38</sup> | 2021 | SVM | Seq. | 0.125 | 0.208 | 0.089 | 0.174 | 0.348 |
| amPEPpy 1.0<sup>39</sup> | 2021 | RF | Seq. | 0.025 | 0.013 | 0.362 | 0.301 | 0.291 |
| Ma et al.<sup>21</sup> | 2022 | BERT, Attention, LSTM | Seq. | 0.174 | 0.786 | 0.098 | 0.182 | 0.358 |
| AMPLify-balanced<sup>40</sup> | 2022 | Bi-LSTM, Attention | Seq. | 0.071 | 0.061 | 0.086 | 0.133 | 0.182 |
| AMPLify-imbalanced<sup>41</sup> | 2023 |  |  | 0.062 | 0.450 | 0.034 | 0.060 | 0.082 |
| iAMPCN<sup>42</sup> | 2023 | CNN | Seq. | 0.130 | 0.498 | 0.074 | 0.139 | 0.254 |
| AMP-BERT<sup>22</sup> | 2023 | BERT | Seq. | 0.264 | 0.200 | 0.390 | 0.577 | **0.731** |
| PGAT-ABPp<sup>24</sup> | 2024 | ProtTrans, GAT | Seq. + Stru. | 0.096 | 0.163 | 0.068 | 0.092 | 0.126 |
| deepAMPNet<sup>23</sup> | 2024 | Bi-LSTM, GCN | Seq. + Stru. | 0.195 | 0.276 | 0.151 | 0.295 | 0.511 |
| SAMP<sup>43</sup> | 2024 | Ensemble RP | Seq. | 0.027 | 0.014 | 0.575 | 0.378 | 0.422 |
| Our approaches | 2024 | ESM-2 (Seq+Tok-CLS) | Seq. | 0.738 | 0.860 | 0.646 | 0.489 | 0.572 |
|  |  | ESM-2 (Tok-CLS) | Seq. | 0.876 | **0.977** | 0.794 | 0.479 | 0.576 |
|  |  | ESM-2 (Tok-CLS-LoRA) | Seq. | **0.900** | 0.972 | **0.838** | 0.590 | 0.669 |

For residue level, which involves classifying individual residues as AMP or non-AMP, was not supported by most tested approaches, including Ma et al. (2022). Therefore, we focused on comparing three variants of our AMP discovery model **(Fig. 2e)**. The results show the end-to-end token-classification models achieved higher precision than the two-step approach, reaching a precision of 0.9 on the testing dataset (**Supplementary Table 2**). The fully fine-tuned and LoRA-fine-tuned token-classification models exhibited similar F1 scores, with the fully fine-tuned model showing slightly higher precision, while the LoRA-fine-tuned model demonstrated slightly better recall (**Fig. 2e**).

These findings highlight the robustness of our framework for both protein-level and residue-level AMP discovery, providing a versatile tool for identifying AMPs from diverse sources.

### 3 AMP-SEMiner predicts ∼1.6M novel candidate AMPs from several habits

Comprehensive benchmarking demonstrates the robustness of the AMP-SEMiner discovery model, ensuring reliable AMP identification. Using this framework, we identified 6,046,125 candidate AMPs (c_AMPs) from diverse microbial proteins across human, non-human animal, and environmental metagenomic datasets (**Supplementary Data 1**). After removing duplicates within the same habitat, 1,670,600 unique c_AMPs were retained (**Fig. 3a**), with some shared across habitats (**Fig. 3b**). Notably, 98.2% of the c_AMPs were shorter than 30 amino acids, slightly shorter than those in other databases. Among all the unique c_AMPs, 8,153 is originated from smORFs (cov >= 0.9). Sequence homology analysis against known AMP databases (APD, dbAMP) identified 146,501 c_AMPs with significant similarity (pident >= 0.9, qcov >= 0.8, mismatch ratio ≤0.1). Remarkably, 91.2% of the c_AMPs were novel (**Supplementary Data 2**).

**Fig. 3.**
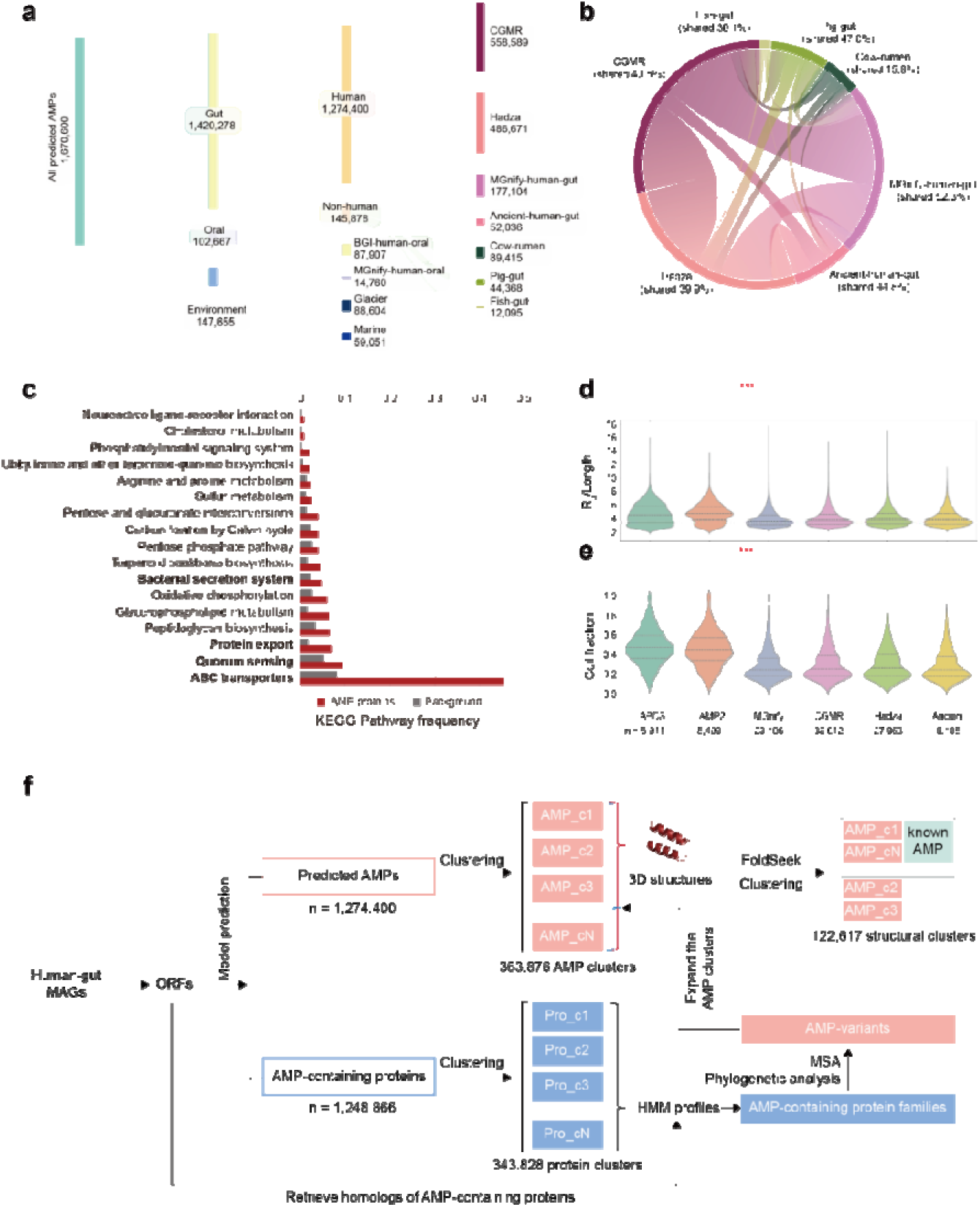
Overview of ∼1.6M predicted AMPs from several habits using AMP-SEMiner discovery model. **(a)** Sankey diagram depicting the distribution of approximately 1.6 million predicted AMPs across various habitats. **(b)** Chord diagram illustrating the interrelationships between predicted AMPs from different habitats. **(c)** KEGG pathwa enrichment analysis highlighting the functional categories associated with AMP-containing proteins, such as secretion, bacterial interaction, and metabolism, compared to background proteins. **(d)** Radius of gyration (Rg) normalized by protein length (Rg/Length) for AMP-containing proteins across different datasets, with *** indicating significant differences. **(e)** Coil fraction of AMP-containing proteins across various datasets, determined using a simplified DSSP algorithm, with *** indicating significant differences. **(f)** Processing of AMPs and AMP-containing proteins int evolutionary context. The ORFs from 4 human gut MAGs are fed into AMP-SEMiner model to obtain 1,274,400 AMP (remove identity sequence) and 1,248,866 AMP-containing proteins (remove identity sequence). And then CD-HIT was used to group the AMP/proteins into 363,876/343,838 clusters. For AMP, we also use FoldSeek to group the 3D of sequence representation and known AMPs into 122,617 structure clusters. To recover the AMP variants overlooked by our model, we use the protein sequences within cluster to build the HMM model to profile ORFs again

Physicochemical analysis showed that these c_AMPs shared characteristics with validated antimicrobial peptides, including a positive net charge, high isoelectric point, amphiphilicity, and the potential to bind membranes or proteins (Boman index) (**Supplementary Fig. 2**). KEGG pathway enrichment analysis showed significant overrepresentation of pathways related to AMP secretion, such as ‘ABC transporters’, and ‘Bacterial secretion system’ **(Fig. 3c)**, consistent with findings from previous study^44^. Structural properties further revealed distinct features compared to known AMP datasets, reinforcing the uniqueness and high quality of these c_AMPs **(Fig. 3d**, **Fig. 3e and Supplementary Fig. 3)**.

To put c_AMPs from four human gut MAGs in an evolutionary context, we identified 363,876/343,828 families by clustering over 1.2M AMP/AMP-containing proteins (**Fig. 3f**). Structural clustering based on AlphaFold2 predictions resulted in 122,617 structural clusters, suggesting a more conservative structure space compared to sequence space **(Fig. 3f)**. Detail of statics on these clusters can be found in **Supplementary Data 3.**

We integrated all sequence and 3D structure data of representative c_AMPs into the online platform MAG-AMPome (https://mag-ampome.aigene.org.cn). This platform enables users to access peptide/AMP sequences, protein annotations, and biochemical properties. Additionally, it provides 3D structures for representative c_AMP sequences. MAG-AMPome aims to facilitate the discovery and application of AMPs, advancing antimicrobial research.

### 4 Experimental results demonstrate the efficiency of AMP-SEMiner

#### 4.1 Experimental validation of several AMPs in vitro assays, with 13 AMPs exhibiting superior efficacy compared to antibiotics

To evaluate the antimicrobial activity of c_AMPs, we selected representatives from 20 clusters based on cluster size and peptide physicochemical properties. Their minimum inhibitory concentration (MIC) values were tested against four pathogenic strains: Gram-negative bacteria, *Escherichia coli* (*E. coli*) and *Pseudomonas aeruginosa* (*P. aeruginosa*), and Gram-positive bacteria, *Staphylococcus aureus* (*S. aureus*) and *Enterococcus faecalis* (*E. faecalis*). As shown in **Fig. 4a**, 9 out of the 20 c_AMPs exhibited antimicrobial activity, completely inhibiting the growth of at least one pathogen at a concentration criterion of 64 μM. The lowest MIC values among these were 4.06 μM and 2.64 μM, respectively. Notably, 5 of the 9 active AMPs demonstrated dual antibiotic activity against both Gram-negative and Gram-positive bacteria, with MIC values (2.66 - 10.54 μM) comparable to or better than at least one of the antibiotics **(Fig. 4a)**. Specifically, the MIC of p9 against *P. aeruginosa* matched that of Polymyxin B, while the MIC of p17 against *E. faecalis* surpassed that of Daptomycin. Details on experimental MIC values of the 20 c_AMPs are provided in **Supplementary Data 4**. Furthermore, eight variants from p17 and one from p1 also demonstrated antimicrobial activity superior to that of antibiotics (**Supplementary Data 5**). In total, 62% (18 out of 29) of all tested c_AMPs were confirmed to be antimicrobial, demonstrating the high efficiency of the AMP-SEMiner model.

**Fig. 4.**
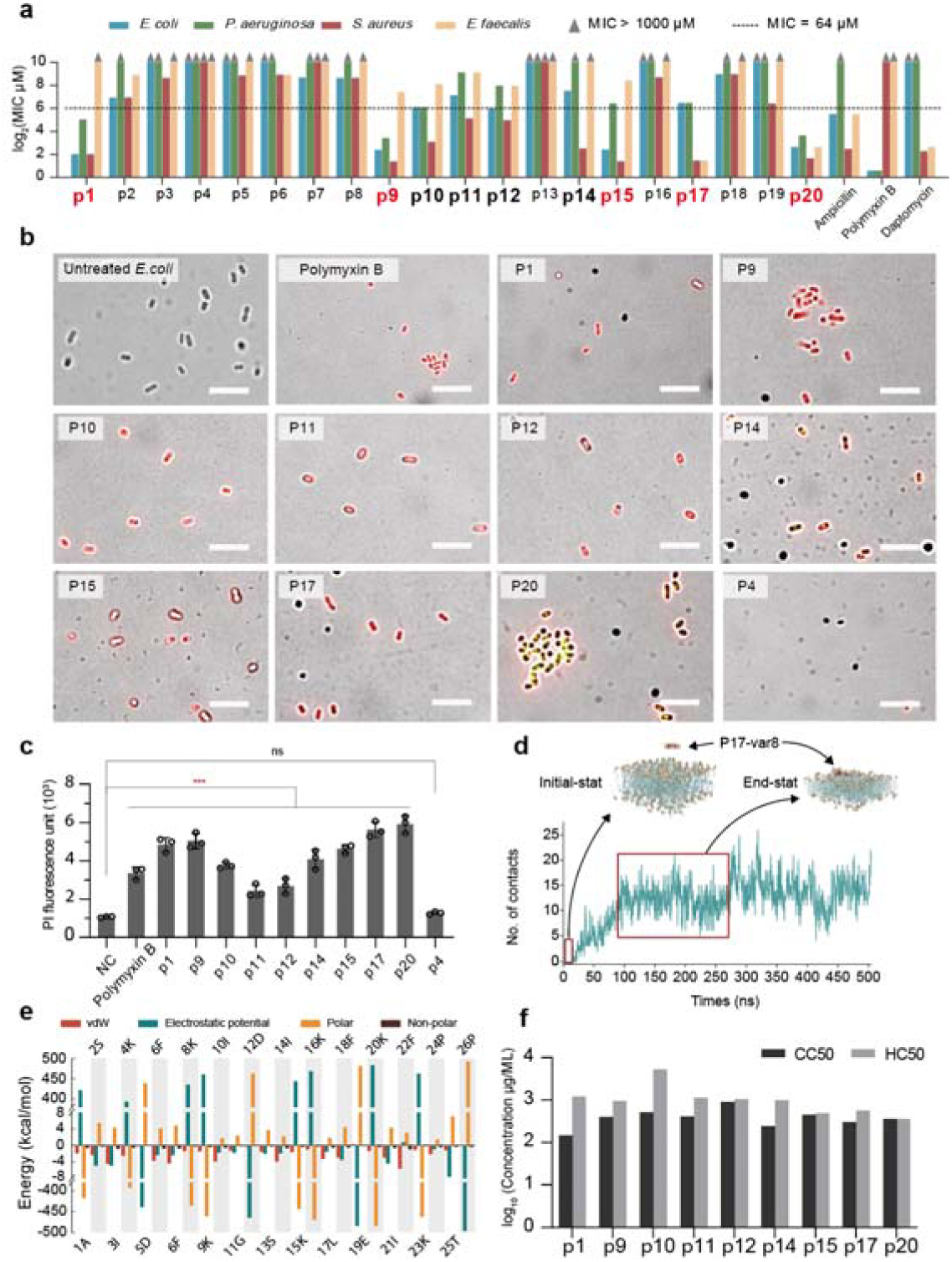
Antimicrobial activity, mechanism of action, cytotoxicity, and hemolytic assessment of AMP candidates, validating the effectiveness of AMP-SEMiner. **(a)** MIC assay results for AMP candidates against four bacterial strains: *E. coli* (ATCC 25922), *S. aureus* (ATCC 25923), *E. faecium* (ATCC 29212), and *P. aeruginosa* (ATCC 27853). Grey triangles represent MIC values greater than 1000 µM, and the dashed horizontal line indicates the 64 µM threshold (antimicrobial concentration criterion). Red-highlighted peptides exhibit superior antimicrobial activity compared to antibiotics; bolded peptides show significant antimicrobial efficacy (MIC ≤ 64 µM). The result show that p1, p9, p10, p11, p12, p14, p15, p17, and p20 exhibit antimicrobial activity. The antimicrobial activity of p1, p9, p15, p17, and p20 surpasses that of antibiotics. **(b)** Phase-contrast microscopy of *E. coli* cells untreated or treated with 9 high active AMPs, 1 non-AMP peptide, and Polymyxin B, and then stained with propidium iodide (red). **(c)** Fluorescence measurement of propidium iodide with *E. coli* (ATCC 25922) cells untreated (control) or treated with AMPs. P-values are for unpaired t-test of each AMP compared to untreated *E. coli* control. Polymyxin B was used as positive control and peptide p4 was a candidate control with no anti-bacterial activity. Scale bar = 5 μm. **(d)** A 500-ns MD simulation of AMP_c423 candidate (p17-var8) interacting with a Gram-positive bacterial membrane (G+PM). A contact is defined as any heavy atom in the AMP being within 3Å of the membrane. The interaction resulted in stable peptide-membrane structures, demonstrating the antimicrobial mechanism as bacterial membrane disruption. **(e)** Molecular mechanics/generalized Born surface area (MM/GBSA) binding free energy calculations for AMP_c423 candidate (p17), demonstrating the role of amino acid residues in peptide-membrane interactions. **(f)** Results of cytotoxicity (CC50), and hemolytic activity (HC50) of AMP candidates. The tested AMP candidates were selected based their high antimicrobial activity in the MIC assays.

#### 4.2 PI staining and molecular simulations reveal AMP mechanisms

Disruption of the bacterial membrane is a key mechanism by AMPs exert their activity. To investigate this, we analyzed peptides with confirmed antimicrobial activity using PI staining (**see Methods**). The results revealed that the red fluorescence readings of tested peptide **(Fig. 4b)** were consistent with their MIC findings, indicating that peptide treatment disrupted the bacterial membrane compared to untreated *E. coli* **(Fig. 4c)**. In contrast, peptide p4, which lacked antimicrobial activity, showed fluorescence levels similar to the negative control, with no detectable red fluorescence, resembling untreated *E. coli*. To further elucidate the mechanism of action, we employed MD simulations and free energy calculations to model the membrane disruption process. Our results confirmed two distinct stages **(Fig. 4d and Supplementary Fig. 4**) consistent with previous study^45^: (1) In the first stage, the peptide adopts a favorable orientation for membrane contact through residue-specific interactions **(Fig. 4d and Supplementary Fig. 4**). (2) The second stage involves peptide insertion, which depends on the interplay among the AMP, membrane, and surrounding solvent (**Fig. 4e and Supplementary Fig. 5**). These findings demonstrate that AMPs disrupt bacterial membranes through a two-stage process, providing mechanistic insights into their antimicrobial activity, highlighting the potential of these peptides as effective antimicrobial agents.

#### 4.3 Active AMP candidates with low cytotoxicity and hemolysis

For AMPs to be effective as antibiotic alternatives, they must exhibit strong antimicrobial activity while minimizing hemolytic effects. Comparative analyses of CC50 and HC50 demonstrated favorable selectivity profiles for peptides with antibacterial activity **(Fig. 4f and Supplementary Fig. 6)**. Most of these peptides had HC50 values exceeding 1000 µM, with a few exceptions ranging between 357.8 and 977 µM **(Supplementary Fig. 7**). Interestingly, peptides with lower MIC values tended to exhibit relatively lower CC50 values, highlighting the trade-off between antimicrobial potency and cytotoxicity. These findings suggest that highly active peptides exert their antimicrobial effects by disrupting bacterial cell membranes, while their low cytotoxicity and hemolytic activity underscore their potential for further development as antimicrobial therapeutics.

### 5 AMP-SEMiner can identify AMPs derived from both smORF and EPs potentially generated within gene protocluster

We introduce a residue-level strategy that enables the detection of AMPs derived from both smORFs and EPs. To validate our method, we constructed several test datasets using AMPs from the APD3 and dbAMP2 databases. Benchmarking results confirm its ability to identify AMPs from both smORFs (**Supplementary Table 8**) and EPs (**Table 1**). A detailed analysis of the AMP_c423 family illustrates the model’s effectiveness. This family includes 82 sequences from four human gut microbiomes, as well as the pig gut microbiome, all validated for antimicrobial activity **(p17 in Fig. 4a)**. Functional annotation revealed that AMP_c423 is encoded within an uncharacterized protein. Genomic context analysis identified conserved upstream and downstream regions flanking AMP_c423 encoded protein, consistently associated with genes encoding Coenzyme PQQ Synthesis Protein D (PqqD) and Cytoplasmic serine protease Peptidase_S24 **(Fig. 5a)**. Peptidase_S24, a serine protease, is known to cleave N-terminal signal peptides^46^, facilitating the secretion of mature peptides **(Fig. 5b)**. SignalP^47^ predictions confirmed the presence of signal peptides in 74 of the 82 sequences **(Supplementary Fig. 8 and Supplementary Data 8)**. Additionally, antiSMASH^48^ classified this family as a gene RRE-containing protocluster due to the presentation of Stand_Alone_Lasso_RRE **(Fig. 5a)**, an enzyme involved in synthesis of lasso peptide with antimicrobial properties^49^.

**Fig. 5.**
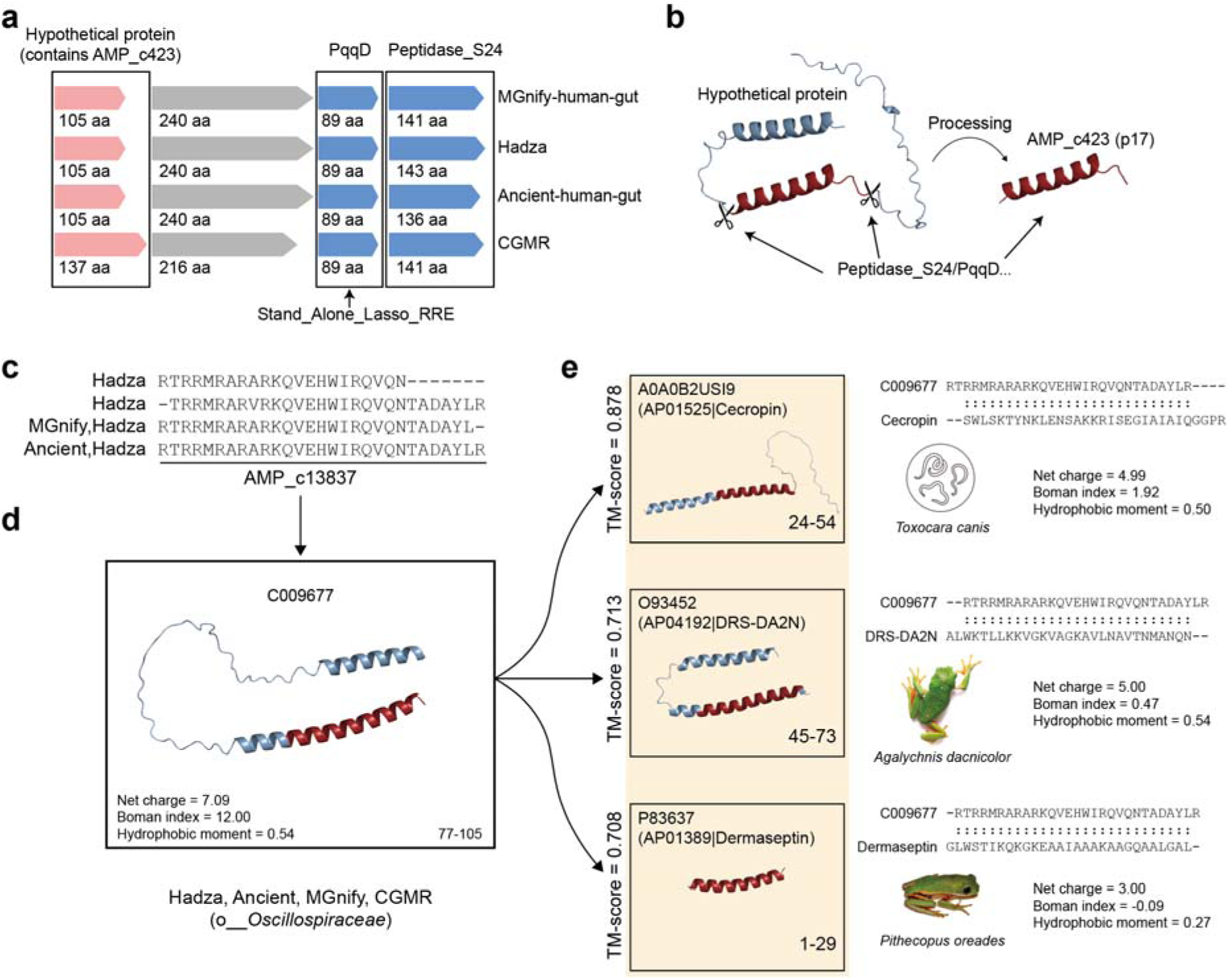
Encrypted AMPs identified by AMP-SEMiner. **(a)** Conserved gene arrangements identified in representatives of AMP_c423-containing hypothetical proteins across four human-gut microbiome datasets. **(b)** Proposed processing mechanism of AMP_c423, potentially involving Peptidase_S24/PqqD catalysis. **(c)** Multiple sequence alignment (MSA) of peptides within the AMP cluster AMP_c13837. **(d)** Structural clustering revealing structural analogs of AMP_c13837 from APD3. **(e)** Three known AMPs from the APD3 database, all originating from eukaryotes, were identified as structural analogs of AMP_c13837.

We also identified AMP_c13837 as an encrypted AMP **(p3 in Fig. 4a)**. This AMP-containing protein family, C009677, was found across all four human gut MAGs **(Fig. 5c and Fig. 5d)**. Furthermore, three well-characterized AMPs from the APD3 database, Cecropin (AP01525) from *Toxocara canis*, DRS-DA2N (AP04192) from *Agalychnis dacnicolor*, and Dermaseptin (AP01389) from *Pithecopus oreades*, were structurally clustered with AMP_c13837, all achieving TM-scores above 0.7 **(Fig. 5e)**.

In total, 7 additional validated c_AMPs **(Supplementary Data 6)** were identified as encrypted AMP. These findings highlight the versatility of our method in overcoming the limitations of previous approaches, which typically detect either encrypted or smORF-derived AMPs, thereby significantly expanding the scope of AMP discovery.

### 6 Evolutionary analysis reveals both the conserved and adaptive evolution of the AMPome in the human gut microbiomes

Changes in microbial community composition are linked to various human diseases^50^, potentially driven by the AMPome of the human gut microbiome^51^. AMPs are vital in host-microbe coevolution, with host-derived AMPs evolving in response to environmental changes. Hence, studying microbial-derived AMPs offers insights into host-microbiota interactions that maintain microbial balance^52^. Using AMP-SEMiner, we identified AMPs produced by four human gut microbes and analyzed their host-specific differences. We observed smaller differences in the composition among three modern human gut microbiomes, whereas ancient human gut microbiomes showed greater variation, with a significant proportion of the microbiota remaining uncharacterized **(Fig. 6a)**. This suggests that the AMPs produced by human gut microbes have evolved in response to changes in their host’s environment.

**Fig. 6.**
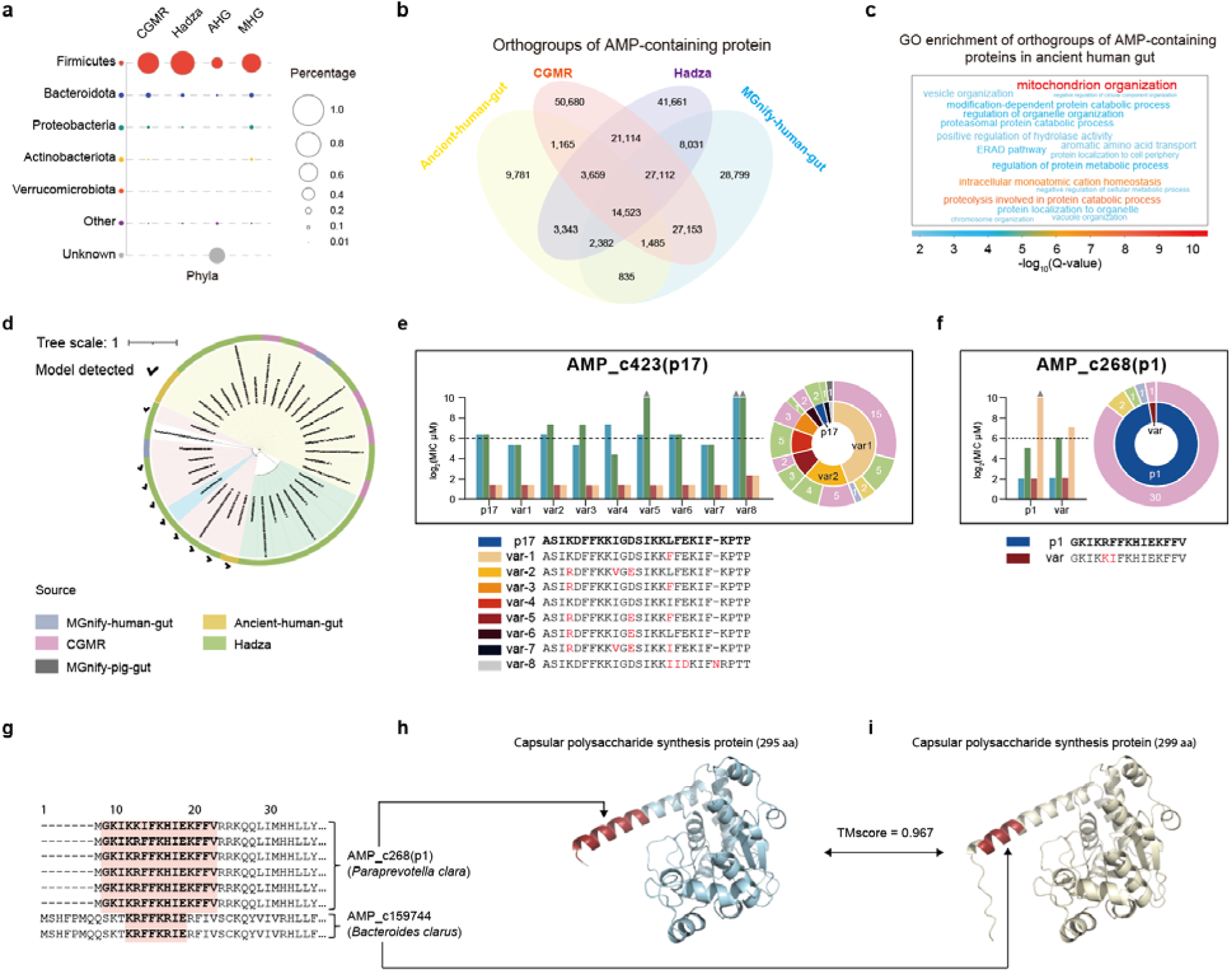
Conserved and adaptive evolution of the AMPome in human gut microbiomes. **(a)** Lineage frequency of AMP-containing genes, categorized at the phylum level. AHG refers to ancient human gut microbiomes, while MHG represents modern human gut microbiomes from the MGnify database. **(b)** Venn diagram illustrating the overlap of AMP-containing gene orthogroups across four human gut microbiomes. **(c)** GO enrichment analysis of AMP-containing orthogroups identified in ancient human gut microbiomes. **(d)** Phylogenetic tree of AMP_c13837 variant peptides, demonstrating how HMM-based searches can recover AMP members previously overlooked by the AMP-SEminer discovery model. **(e-f)** MIC assay results for peptide variants p1 and p17. Hierarchical pie charts display the distribution of peptide variants (inner ring) and their respective source datasets (outer ring). **(g)** Multiple sequence alignment of the N-terminal regions of AMP-containing proteins from two AMP clusters, AMP_c268 and AMP_c159744. Highlighted regions indicate predicted AMPs. These proteins originate from *Paraprevotella clara* and *Bacteroides clarus*, respectively. **(h)** Predicted 3D structure of the AMP_c268-containing protein. **(i)** Predicted 3D structure of the AMP_c159744-containing protein. **(h)** and **(i)** share highly similar 3D structures (TM-score = 0.967), with differences in the N-terminal region.

To examine variations in AMP-encoding genes, we constructed orthogroups (OGs) based on protein sequences. The analysis revealed that these human gut microbes share 14,523 AMP-containing Ogs while retaining unique OGs specific to each human gut microbes **(Fig. 6b).** Shared OG families were linked to transporters, biosynthesis, metabolism and quorum sensing **(Supplementary Data 7)**, while unique OGs exhibited distinct function (**Fig. 6c and Supplementary Fig. 9**). These findings indicate that AMP-encoding genes in four human gut microbiomes exhibit both evolutionary conservation and adaptation, aligning with previous observations in specific environmental contexts^44^.

To further investigate the evolution of the human gut AMPome, we clustered the AMP-containing proteins into 343,828 families (**Fig. 3f**) and constructed HMM profile for each family. These profiles were then used to search for additional homologs in MAGs, allowing us to recover sequences previously missed by the AMP-SEMiner discovery model, thereby improving the robustness of the method (**Fig. 6d**).

To investigate the characteristics of conserved AMP families, we analyzed the AMP_c423 family, consisting of 82 members with 9 distinct sequence variants. Given their very similar 3D structures, these variants showed comparable antimicrobial activity in vitro (**Fig. 6e and Supplementary Data 5)**. Notably, mutations within these variants followed specific patterns: substitutions between charged residues (e.g., K→R, E→D) or between hydrophobic residues (e.g., I→V, L→F, I→F) generally preserved antimicrobial function. These mutations maintained essential physicochemical properties, such as charge or hydrophobicity, which are critical for activity. MD simulations further confirmed that these mutations had minimal impact on peptide-membrane interaction energy **(Fig. 4e and Supplementary Fig. 5)**. Although there are slight sequence variations within these families, the antimicrobial efficacy and mechanisms of the peptides remain unchanged. This example illustrates that, under natural conditions, conserved AMP families can tolerate certain mutations without completely losing their antimicrobial function, in contrast to the loss of activity observed in AMP produced by *Drosophila*^53^. Another validated example is AMP_c9875 also showed conserved evolutionary across gut microbiomes (**Supplementary Data 6**).

To explore the adaptive evolution of AMP families, we analyzed AMP_c268 **(Fig. 6f)**, which consists of 35 AMP candidates within a protein family annotated as “capsular polysaccharide synthesis protein.” Of these, 34 were identical to peptide p1, with one variant showing comparable MIC values. This protein family also includes a related AMP family, AMP_c159744, comprising proteins from *Paraprevotella clara* and *Bacteroides clarus*. Both microbes, members of the *Bacteroidaceae* family, represent closely related but functionally distinct lineages: *Paraprevotella clara* is phylogenetically and phenotypically most similar to *Prevotella* species which is specializes in fiber degradation in plant-rich diets^54, 55^, while *Bacteroides clarus* is a generalist with broader metabolic capabilities^56^. These differences highlight the adaptive responses of gut microbes to dietary and environmental changes, contributing to host health. Additionally, six other AMP clusters exhibited evidence of adaptive evolution (**Supplementary Data 6**). Together, these findings reveal both conserved and adaptive evolutionary patterns of AMPs in gut microbiota, offering valuable insights into their functional roles and evolutionary trajectories.

## Discussion

The discovery of natural AMPs is vital for addressing the growing challenge of antibiotic resistance. We developed an AMP discovery model capable of identifying both smORF-derived and EP-derived AMPs, utilizing a residue-level classifier built on the advanced ESM-2. Benchmarking demonstrated its superiority over existing methods at the protein and residue level. We extended this model into AMP-SEMiner, a robust computational framework that integrates structural clustering and evolutionary analysis to identify AMP clusters and uncover AMP families with natural variants. Using AMP-SEMiner, we predicted over 1 million AMPs from metagenomic datasets. Experimental validation of 29 candidates, confirmed antimicrobial activity in 18 cases. These findings highlight the efficiency and robustness of our AMP discovery model and AMP-SEMiner framework, significantly broadening the scope of AMP discovery and demonstrating the potential of computational methods to accelerate the development of novel antimicrobials.

Key insights from our AMP-SEMiner exploration include:

1. Cross-kingdom similarity. AMPs derived from microbial proteins, such as AMP_c13837, exhibited high structural similarity to known eukaryotic AMPs, emphasizing the potential of microbial-origin proteins for AMP discovery.
2. Environmental influence on AMP evolution. Some AMPs, like AMP_c423, were shared across human gut microbiomes, while others, like AMP_c159744 (e.g., expanded in gut microbes from the modern Chinese cohort), were cohort-specific, suggesting that environmental factors such as diet may shape the evolutionary trajectory of AMPs.
3. Unannotated proteins with antimicrobial roles. Among validated AMPs, we identified previously unannotated hypothetical proteins, like AMP_c423, with antimicrobial activity. Genomic context exploration provided insights into this protein may related to the other protein, while PI staining and MD simulations revealed their antimicrobial mechanism as bacterial membrane disruption.

Limitations of this study include the following: (1) Bias toward shorter AMPs. Filtering criteria focused on fragments longer than 5 amino acids, limiting the discovery of longer peptides, which is acceptable for current AMP applications (**Supplementary Fig. 2**). (2) Trade-off between potency and safety. The natural AMPs discovered exhibited a balance between antimicrobial potency and safety, likely reflecting the equilibrium within human microbiota. (3) Limited exploration of mechanisms. While MD simulations and PI staining revealed bacterial membrane disruption as a key mechanism, other potential mechanisms remain unexplored, warranting further investigation.

Subsequent potential studies based on our results include the following: (1) Expanding the AMP-SEMiner framework to identify longer AMPs and refine residue-level classifiers for broader peptide discovery. (2) Investigating additional mechanisms of AMP action beyond bacterial membrane disruption, such as targeting intracellular components. (3) Exploring environmental factors influencing AMP evolution and their implications for personalized medicine. (4) Developing synthetic AMPs based on natural templates to optimize antimicrobial activity while minimizing cytotoxicity.

## Methods

### 1. Development of AMP-SEMiner for mining AMP from MAGs

#### 1.1 Construction of training, validation and testing dataset using known AMPs

Experimentally validated AMP sequences were obtained from the Giant Repository of AMP Activities (GRAMPA) dataset^34^, combining data from dbAMP^35^ (v2.0). After removing duplicates, the non-redundant AMP sequences were aligned to the UniProt database^57^ using blastp^58^ (v2.15.0). For AMP sequences with lengths <= 30 amino acids (aa), the “-task short” parameter was used. Hits were filtered based on the following criteria:1) Proteins longer than 300 aa were excluded. 2) Alignments with no gaps were retained. 3) Mismatches were limited to no more than 2 aa for length >10 aa or 1 aa for length <= 10 aa. The remaining well-aligned AMP-hit proteins were randomly split into training, validation, and testing datasets in a 7:2:1 ratio. For the negative dataset, we employed the same strategy as described in Ref^42^. Proteins were retrieved on January 22, 2024, using the query term: “(cc_scl_term:SL-0086) NOT antimicrobial NOT antibiotic NOT antiviral NOT antifungal NOT effector NOT excreted AND (length:[* TO 300])”. Redundant sequences with >= 90% similarity were removed using CD-HIT^59^. Proteins in the negative dataset were further filtered to ensure sequence similarity was < 90% with those in the training dataset. The resulting non-redundant proteins were randomly split into training, validation, and testing datasets at a 7:2:1 ratio, serving as AMP-negative data.

#### 1.2 Construction of independent testing datasets

Since GRAMPA includes older versions of APD3 and LAMP2 data, we exclusively used the updated datasets, APD (2024) and LAMP2 (2020), as independent testing sets. To ensure fairness, redundant AMP sequences with >= 90% similarity were removed using CD-HIT.

#### 1.3 Fine-tuning ESM-2 for residue-level AMP identification

To identify both smORF and EP AMPs, we fine-tuned the pretrained ESM-2 model (650M) for a token classification task. This task involved classifying each residue in a protein as AMP-positive or AMP-negative. For implementation, we utilized the EsmForTokenClassification and EsmForSequenceClassification classes provided by HuggingFace (https://github.com/huggingface/transformers) for residue-level and protein-level identification tasks, respectively. Three training approaches were explored:

1. 2-Step model: the AMP sequence classification model and the token classification model were trained independently.
2. Fully fine-tuned model: all parameters of the ESM-2 model and the fully connected layers within EsmForTokenClassification were trained simultaneously.
3. LoRA fine-tuning model: the LoRA (Low-Rank Adaptation) method was implemented using the PEFT (Parameter-Efficient Fine-Tuning) library (https://huggingface.co/docs/peft/en/index), with a LoRA rank set to 8^60^.

#### 1.4 Benchmarking and model performance evaluation

A total of 12 published tools were included in the benchmarking analysis: Macrel^20^, AmpGram^37^, ampir^38^, amPEPpy 1.0^39^, Ma-AMPminer^21^, AMPlify (balanced^40^ and imbalanced^41^), iAMPCN^42^, AMP-BERT^22^, PGAT-ABPp^24^, deepAMPNet^23^ and SAMP^43^. Alongside these, we evaluated our AMP discovery models, including the 2-Step model, the fully fine-tuned model, and the LoRA fine-tuned model, using both the testing and independent datasets (APD3 and LAMP2).

1. Protein-level Evaluation. For the 12 published tools, protein predictions were classified as AMP or non-AMP. In our AMP discovery models, a protein was defined as AMP-containing if it included more than 5 consecutive residues predicted to be AMP-positive. Proteins not meeting this criterion were classified as AMP-negative.
2. Residue-level Evaluation. None of the 12 published tools supported residue-level evaluation. Our AMP discovery models, however, were assessed at the residue level, enabling per-residue analysis.

Model performance was evaluated using the Evaluate library (https://huggingface.co/docs/evaluate), with metrics such as precision, recall, and F1-score calculated to provide a comprehensive performance assessment.

#### 1.5 MAGs data collection

Prokaryotic MAGs and metagenomic sequencing data used for AMP detection were sourced from various public datasets. These included the CGMR38 dataset^61^ (https://ngdc.cncb.ac.cn/bioproject/browse/PRJCA017330), Hadza dataset^62^, 4D-SZ human oral dataset^63^, ancient human gut dataset^64^, Glacier Microbiomes dataset^65^, and MGnify datasets^66^: human-gut-v2.0, human-oral-v1.0, cow-rumen-v1.0, pig-gut-v1.0, fish-gut-v1.0, zebrafish-fecal-v1.0, and marine-v1.0 (https://www.ebi.ac.uk/metagenomics/browse/genomes).

For the ancient human gut dataset, raw sequencing data from eight ancient human gut microbiomes were downloaded from the Sequence Read Archive (SRA) under BioProject accession number PRJNA561510, as the assembled MAGs were not publicly available. We adhered to the same read processing and assembly pipeline as described in the original study^64^. First, AdapterRemoval^67^ was used to trim adapter sequences from the reads, followed by KneadData (https://github.com/biobakery/kneaddata) to remove contamination from host. Cutadapt^68^ was then applied to ensure the complete removal of any remaining adapters, resulting in a set of clean reads. These clean reads were assembled into MAGs using MEGAHIT^69^ with default parameters. Bowtie2^70^ was used to map the clean reads to the MAGs, and Samtools^71^ was employed to generate sorted BAM files. MetaBAT2^72^ was subsequently utilized to bin the contigs, and taxonomy classification was conducted using GTDB-Tk with the GTDB database^73^.

#### 1.6 AMP candidate discovery

ORFs were predicted using pyrodigal^74^ v.3.5.2 (https://github.com/althonos/pyrodigal). The parameters ‘-- max-overlap’ and ‘--min-gene’ were both set to 15 to ensure identification of all ORFs with a minimum length of 5 amino acids (aa). ORFs with lengths <= 300 aa were selected as input protein sequences for AMP identification using our fully fine-tuned AMP discovery model. Protein fragments and smORFs composed of more than five consecutive residues were classified as AMP candidates.

#### 1.7 Orthogroups analysis

OGs were inferred using OrthoFinder with the “-s” parameter set to MMseqs^75^ for AMP-containing proteins derived from four human gut MAGs. Gene Ontology (GO) annotations were obtained using eggNOG-mapper^76^. Following the approach used in previous study^77^, GO terms for each OG were converted to level 5, redundant GO terms within the same OG were removed. Using the shared OGs as a background, enrichment analysis was conducted to identify unique functional annotations across the four human gut microbiomes.

### 2. Structural analysis of AMP candidates

AMP candidates and AMP-containing proteins were clustered using CD-HIT, with the ‘-c’ parameter set to 0.95 for AMP candidates and 0.8 for AMP-containing proteins. The three-dimensional (3D) structures of the representative sequences from each AMP-containing protein cluster were predicted using ColabFold^78^ v1.5.5: AlphaFold2 using MMseq2 (https://github.com/sokrypton/ColabFold/blob/main/AlphaFold2.ipynb). The AMP fragments within these proteins were extracted using biopython^79^ v.1.79. Structural clustering was performed separately for AMPs and AMP-containing proteins using FoldSeek^31^ easy-cluster with parameters ‘--tmscore-threshold 0.5 --lddt-threshold 0.6 --alignment-type 1 --cluster-mode 1’. For AMP structure clustering, known AMP structures from the APD3 database were also included in the analysis to provide a comprehensive comparison. Pairwise structural similarity was calculated using USalign^80^. The calculation of structural descriptors, including radii of gyration and coil fractions, was performed as described in a previous study^81^ (https://github.com/qianyuantang/stat-trend-protein-evo). The calculation of sequence physicochemical properties was performed biopython and peptides tool (https://github.com/althonos/peptides.py).

### 3. Evolutionary analysis of AMP clusters

Hidden Markov Model (HMM) profiles for clusters of AMP-containing proteins were generated using HMMER^33^. These profiles were employed to search all microbial ORFs to identify potential homologs, forming protein families. Functional annotations, including KEGG pathway annotations, for the protein families were performed using eggNOG-mapper^76^ v2 (http://eggnog-mapper.embl.de/). Multiple sequence alignments (MSAs) for each AMP-containing protein family were constructed using MAFFT^82^ v7.508 with 1000 iterations. Phylogenetic trees were then inferred using IQ-TREE^83^ v1.6.12, employing automatic model selection and 1000 bootstrap replicates. The resulting trees were visualized using iTOL^84^ v5 (https://itol.embl.de/itol.cgi). All variant peptides identified from the MSAs of a specific AMP-containing protein family were defined as an AMP family. These AMP families were subsequently subjected to experimental validation.

### 4. Experimental validation of AMP

#### 4.1 Measurement of minimum inhibitory concentration

All peptides used in the experiments were synthesized by GenScript (Suzhou, China). The MIC assay of the designed AMPs was performed using the broth microdilution method according to the reported protocols^85^. The bacterial strains used in this study were *E. coli* (ATCC 25922), *S. aureus* (ATCC 25923), *E. faecium* (ATCC 29212), and *P. aeruginosa* (ATCC 27853). The synthesized peptides were dissolved in 1% DMSO at a concentration of 10 mg/mL for storage (10×). Briefly, to prepare the solution for the MIC assay, the dissolved peptide solutions were subjected to two-fold serial dilutions across columns 1-10 of sterile 96-well plates, with columns 11 and 12 serving as controls containing Mueller-Hinton Broth (MHB). Each dilution (50 µL) was pipetted into the wells before the addition of bacterial suspension. The plates were kept covered when not in use to prevent contamination. Bacterial cultures were grown overnight in MHB with shaking at 37°C, then diluted to 1.0 × 10 CFU/mL with MHB. A 50 µL aliquot of each bacterial suspension was added to columns 1–11 of the 96-well plates, which contained the peptide solutions, while column 12 contained MHB only. The plates were incubated at 37°C for 16-20 h. The MIC values were considered as the highest concentration of the peptides at which no visible bacterial growth was observed. All assays were performed in triplicate to ensure statistical reliability.

#### 4.2 Mechanism of action analysis using propidium iodide

The *E. coli* in the exponential phase was resuspended in PBS and adjusted to an OD600 value of 1.0. A 10 µL aliquot of the bacterial suspension was mixed with 10 µL of AMP at a final concentration of 4× MIC, and then incubated at 37 °C for 1 h. To both AMP-treated and untreated samples, 20 μM of propidium iodide (PI, Solarbio, #C0080) was added and incubated at 37 °C for 30 min away from light. Fluorescence was recorded using the Infinite® Eplex plate reader (Tecan) with excitation at 535 nm and emission at 615 nm. In addition, the reaction mixture was applied to a slide and imaged using the Leica DM4B Upright Microscope equipped with a 100x semi-apochromatic objective.

#### 4.3 Assessment of cytotoxicity

The cytotoxicity assay was performed on IEC-6 intestinal epithelial cells (ATCC, #CRL-1592™) using the CellTiter 96® AQueous One Solution Cell Proliferation Assay (Promega, #G3580). Cells were cultured in media containing DMEM high glucose with 10% fetal bovine serum. Cells were diluted to a density of 1.0 × 104 cells per 100 µL and seeded into 96-well plates for 24 h. Following this incubation, the cells were exposed to a series of peptide concentrations, prepared by two-fold serial dilutions from 1000 to 31.25 μg/mL. A 4 µL aliquot of each dilution was added to the wells, and the plates were incubated at 37 °C with 5% CO₂ for a further 24 h. Then, 20 µL of CellTiter 96® AQueous One Solution was added to each well, and after incubation at 37 °C for 60 min, the absorbance was measured at 490 nm, with corrections made using wells containing medium only. CC50 values, representing the concentration of each AMP that kills 50% of the cells, were determined by fitting a 4-parameter logistic model (4PL) using R 4.3.3.

#### 4.4 Assessment of Hemolytic Activity

Rat erythrocytes (Sbjbio, #SBJ-RBC-RAT004) were washed with phosphate-buffered saline (PBS) and resuspended to approximately 2% in PBS. AMPs were two-fold serial diluted from 1000 to 31.25 μg/mL. Each 20 µL AMP dilution was combined with 180 µL of the 2% erythrocyte suspension. The mixtures were incubated at 37 °C for 1 h after sealing. After centrifugation at 1000 × g for 5 min at room temperature, 50 µL of the supernatant was transferred to a transparent 96-well plate and the absorbance was measured at 540 nm. Triton X-100 treated erythrocytes served as a positive control and the PBS treated erythrocytes acted as a negative control. The HC50 values, representing the concentration of AMP required to lyse 50% of the RBCs, were determined by fitting a 4-parameter logistic model (4PL) using R 4.3.3.

### 5 MD simulations to investigate antimicrobial mechanism

The Gram-positive bacterial membrane (G+PM) was modeled using CHARMM-GUI^86^ with 100 lipid molecules per leaflet and optimized using AmberTools^87^ due to issues with specific lipid parameters. Each modelling system included one AMP molecule placed ∼15 Å from the membrane. Simulations were performed with the Amber^88^ *ff19SB* force field for proteins and *gaff* for ligands, using TIP3P water boxes with a 10 Å buffer. Before MD simulation, the systems underwent energy minimization in two stages: restrained backbone atoms (10 kcal·mol⁻¹·Å⁻² elastic constants) followed by unrestrained minimization for 5,000 steps. MD simulations were conducted using Amber. Systems were heated to 300 K in the NVT ensemble and equilibrated at 310.15 K under NPT conditions. And then A 500 ns production run was performed with SHAKE^89^ constraints and a 2 fs timestep, saving coordinates every 5 ps. Binding free energies were calculated using MM/GBSA, decomposing ΔG_bind into molecular mechanics energy (ΔE_MM), solvation energy (ΔG_sol), and entropy (-TΔS). ΔE_MM included electrostatic (ΔE_ele), van der Waals (ΔE_vdw), and internal energies, while ΔG_sol comprised polar (ΔG_GB) and nonpolar (ΔG_SA) contributions. Entropy changes were neglected due to limited impact on binding accuracy^90^.

### 6 Statistics & Reproducibility

The Mann-Whitney U test was used to assess whether the differences in physicochemical properties between AMP candidates and known AMPs from the database were statistically significant.

Additionally, Fisher’s exact test was applied to evaluate whether the frequencies of proteins occurring in KEGG pathways differed significantly between the foreground datasets and the background datasets. Statistical significance is indicated by asterisks (***), representing P < 0.001.

No statistical method was used to predetermine sample size. No data were excluded from the analyses. The experiments were not randomized. The Investigators were not blinded to allocation during experiments and outcome assessment.

## Data availability

The model weights of AMP-SEMiner and a minimum dataset to run AMP-SEMiner are available in Zenodo in DOI [10.5281/zenodo.14348290]. Source data are provided with this paper. We also provide a web-portal (http://mag-ampome.aigene.org.cn) to access all the AMP candidates.

## Code availability

The custom code of AMP-SEMiner, together with the trained models for mining AMPs, are available at GitHub repositories (https://github.com/zjlab-BioGene/AMP-SEMiner). We also provide a quick example on Colab for demonstration (https://colab.research.google.com/drive/1-O8U7M6UTtSaMQqm3sX7ZOtkUCVEmmLt?usp=sharing).

## Acknowledgements

This work was supported by National Natural Science Foundation of China (Grant No. 82450113) to X.X.H, and “Pioneer” and “Leading Goose” R&D Program of Zhejiang (2024SSYS0007). We acknowledge the technical support provided by the Bio-computation and Validation Facility of Zhejiang Lab. Our gratitude also extended to the Material Scientific Cores of Zhejiang Lab for their technical assistance.

## Author Contributions Statement

J.F.Z., X.X.H. and W.H.L. conceived the project. J.F.Z., W.H.L. and M.H.G. built the AMP-SEMiner framework. W.H.L., J.F.Z., M.H.G., B.C.H., Z.H.Z., X.P.Z., X.Y.J. and L.G. analysed the data. W.H.L. and M.H.G. built and trained the AI models. X.X.H., J.F.Z., B.C.H. and Z.H.Z. designed and performed biochemical experiments. E.C.W. and J.F.Z. designed and performed MD simulation experiments. J.F.Z., W.H.L., B.C.H., E.C.W. and X.X.H. wrote this manuscript. M.H.G., J.F.Z., T.C. and L.Q.F. designed and built the AMP-SEMiner web-portal. Y.B.Y. provided computational resources for generating 3D structures of proteins and AMPs. This manuscript was reviewed and approved by all authors.

## References

1. Chinemerem Nwobodo, D., et al. Antibiotic resistance: The challenges and some emerging strategies for tackling a global menace. 36, e24655 (2022).

2. Huang, K.-Y. et al. Identification of natural antimicrobial peptides from bacteria through metagenomic and metatranscriptomic analysis of high-throughput transcriptome data of Taiwanese oolong teas. BMC systems biology 11, 29–44 (2017).

3. Cotter, P.D., Ross, R.P. & Hill, C. Bacteriocins—a viable alternative to antibiotics? Nature Reviews Microbiology 11, 95–105 (2013).

4. Sohlenkamp, C. & Geiger, O. Bacterial membrane lipids: diversity in structures and pathways. FEMS Microbiol Rev 40, 133–159 (2016).

5. Ghosh, A. et al. Indolicidin targets duplex DNA: structural and mechanistic insight through a combination of spectroscopy and microscopy. ChemMedChem 9, 2052–2058 (2014).

6. Zhao, X. et al. Isolation and identification of antifungal peptides from Bacillus BH072, a novel bacterium isolated from honey. Microbiol Res 168, 598–606 (2013).

7. Luan, X. et al. Cytotoxic and antitumor peptides as novel chemotherapeutics. Natural product reports 38, 7–17 (2021).

8. Torres, M.D.T. et al. Mining for encrypted peptide antibiotics in the human proteome. Nat Biomed Eng 6, 67–75 (2022).

9. Nolan, E.M. & Walsh, C.T. How nature morphs peptide scaffolds into antibiotics. ChemBioChem 10, 34–53 (2009).

10. Singh, N. & Abraham, J. Ribosomally synthesized peptides from natural sources. J Antibiot (Tokyo) 67, 277–289 (2014).

11. Oyama, L.B. et al. The rumen microbiome: an underexplored resource for novel antimicrobial discovery. npj Biofilms and Microbiomes 3, 33 (2017).

12. Lang, H. et al. Identification of peptides from honeybee gut symbionts as potential antimicrobial agents against Melissococcus plutonius. Nature Communications 14, 7650 (2023).

13. Chen, S. et al. The discovery of antimicrobial peptides from the gut microbiome of cockroach *Blattella germanica* using deep learning pipeline. bioRxiv, 2024.2002.2012.580024 (2024).x1

14. King, A.M. et al. Systematic mining of the human microbiome identifies antimicrobial peptides with diverse activity spectra. Nature Microbiology 8, 2420–2434 (2023).

15. Torres, M.D.T. et al. Mining human microbiomes reveals an untapped source of peptide antibiotics. Cell 187, 5453–5467 (2024).

16. Crits-Christoph, A., Diamond, S., Butterfield, C.N., Thomas, B.C. & Banfield, J.F. Novel soil bacteria possess diverse genes for secondary metabolite biosynthesis. Nature 558, 440–444 (2018).

17. Maasch, J., Torres, M.D.T., Melo, M.C.R. & de la Fuente-Nunez, C. Molecular de-extinction of ancient antimicrobial peptides enabled by machine learning. Cell Host Microbe 31, 1260–1274 e1266 (2023).

18. Ostaff, M.J., Stange, E.F. & Wehkamp, J. Antimicrobial peptides and gut microbiota in homeostasis and pathology. EMBO Mol Med 5, 1465–1483 (2013).

19. Alexander, P.J. et al. Microbiome-derived antimicrobial peptides show therapeutic activity against the critically important priority pathogen, Acinetobacter baumannii. npj Biofilms and Microbiomes 10, 92 (2024).

20. Santos-Junior, C.D., Pan, S., Zhao, X.M. & Coelho, L.P. Macrel: antimicrobial peptide screening in genomes and metagenomes. PeerJ 8, e10555 (2020).

21. Ma, Y. et al. Identification of antimicrobial peptides from the human gut microbiome using deep learning. Nature Biotechnology 40, 921–931 (2022).

22. Lee, H., Lee, S., Lee, I. & Nam, H. AMP-BERT: Prediction of antimicrobial peptide function based on a BERT model. Protein Sci 32, e4529 (2023).

23. Zhao, F. et al. deepAMPNet: a novel antimicrobial peptide predictor employing AlphaFold2 predicted structures and a bi-directional long short-term memory protein language model. PeerJ 12, e17729 (2024).

24. Hao, Y., Liu, X., Fu, H., Shao, X. & Cai, W. PGAT-ABPp: harnessing protein language models and graph attention networks for antibacterial peptide identification with remarkable accuracy. Bioinformatics 40 (2024).

25. Chen, X. et al. Roles and mechanisms of human cathelicidin LL-37 in cancer. Cellular Physiology and Biochemistry 47, 1060–1073 (2018).

26. Lin, Z. et al. Evolutionary-scale prediction of atomic-level protein structure with a language model. Science 379, 1123–1130 (2023).

27. Elnaggar, A. et al. ProtTrans: Toward Understanding the Language of Life Through Self-Supervised Learning. IEEE transactions on pattern analysis and machine intelligence 44, 7112–7127 (2022).

28. Li, W. et al. Discovering CRISPR-Cas system with self-processing pre-crRNA capability by foundation models. bioRxiv, 2024.2003.2011.583506 (2024).

29. Yu, T. et al. Enzyme function prediction using contrastive learning. Science 379, 1358–1363 (2023).

30. Jumper, J. et al. Highly accurate protein structure prediction with AlphaFold. Nature 596, 583–589 (2021).

31. van Kempen, M. et al. Fast and accurate protein structure search with Foldseek. Nat Biotechnol 42, 243–246 (2024).

32. Wang, G., Li, X. & Wang, Z. APD3: the antimicrobial peptide database as a tool for research and education. Nucleic Acids Research 44, D1087–D1093 (2016).

33. Johnson, L.S., Eddy, S.R. & Portugaly, E. Hidden Markov model speed heuristic and iterative HMM search procedure. BMC Bioinformatics 11, 431 (2010).

34. Witten, J. & Witten, Z. Deep learning regression model for antimicrobial peptide design. BioRxiv, 692681 (2019).

35. Jhong, J.H. et al. dbAMP 2.0: updated resource for antimicrobial peptides with an enhanced scanning method for genomic and proteomic data. Nucleic Acids Res 50, D460–D470 (2022).

36. Ye, G. et al. LAMP2: a major update of the database linking antimicrobial peptides. Database (Oxford) 2020 (2020).

37. Burdukiewicz, M. et al. Proteomic Screening for Prediction and Design of Antimicrobial Peptides with AmpGram. Int J Mol Sci 21 (2020).

38. Fingerhut, L., Miller, D.J., Strugnell, J.M., Daly, N.L. & Cooke, I.R. ampir: an R package for fast genome-wide prediction of antimicrobial peptides. Bioinformatics 36, 5262–5263 (2021).

39. Lawrence, T.J. et al. amPEPpy 1.0: a portable and accurate antimicrobial peptide prediction tool. Bioinformatics 37, 2058–2060 (2021).

40. Li, C. et al. AMPlify: attentive deep learning model for discovery of novel antimicrobial peptides effective against WHO priority pathogens. BMC Genomics 23, 77 (2022).

41. Li, C., Warren, R.L. & Birol, I. Models and data of AMPlify: a deep learning tool for antimicrobial peptide prediction. BMC Res Notes 16, 11 (2023).

42. Xu, J. et al. iAMPCN: a deep-learning approach for identifying antimicrobial peptides and their functional activities. Briefings in Bioinformatics 24, bbad240 (2023).

43. Feng, J. et al. SAMP: Identifying antimicrobial peptides by an ensemble learning model based on proportionalized split amino acid composition. Brief Funct Genomics 23, 879–890 (2024).

44. Santos-Junior, C.D. et al. Discovery of antimicrobial peptides in the global microbiome with machine learning. Cell 187, 3761–3778 e3716 (2024).

45. Hong, H., Choi, H.K. & Yoon, T.Y. Untangling the complexity of membrane protein folding. Curr Opin Struct Biol 72, 237–247 (2022).

46. Ekici, O.D., Paetzel, M. & Dalbey, R.E. Unconventional serine proteases: variations on the catalytic Ser/His/Asp triad configuration. Protein Sci 17, 2023–2037 (2008).

47. Teufel, F. et al. SignalP 6.0 predicts all five types of signal peptides using protein language models. Nature Biotechnology 40, 1023–1025 (2022).

48. Blin, K. et al. antiSMASH 7.0: new and improved predictions for detection, regulation, chemical structures and visualisation. Nucleic Acids Res 51, W46–W50 (2023).

49. Hegemann, J.D., Zimmermann, M., Xie, X. & Marahiel, M.A. Lasso peptides: an intriguing class of bacterial natural products. Acc Chem Res 48, 1909–1919 (2015).

50. Hooper, L.V. & Gordon, J.I. Commensal host-bacterial relationships in the gut. Science 292, 1115–1118 (2001).

51. Gallardo-Becerra, L., Cervantes-Echeverría, M., Cornejo-Granados, F., Vazquez-Morado, L.E. & Ochoa-Leyva, A. Perspectives in Searching Antimicrobial Peptides (AMPs) Produced by the Microbiota. Microbial Ecology 87, 8 (2023).

52. Ageitos, J.M., Sánchez-Pérez, A., Calo-Mata, P. & Villa, T.G. Antimicrobial peptides (AMPs): Ancient compounds that represent novel weapons in the fight against bacteria. Biochemical pharmacology 133, 117–138 (2017).

53. Hanson, M.A., Grollmus, L. & Lemaitre, B. Ecology-relevant bacteria drive the evolution of host antimicrobial peptides in Drosophila. Science 381, eadg5725 (2023).

54. Ley, R.E. Gut microbiota in 2015: Prevotella in the gut: choose carefully. Nat Rev Gastroenterol Hepatol 13, 69–70 (2016).

55. Morotomi, M., Nagai, F., Sakon, H. & Tanaka, R. Paraprevotella clara gen. nov., sp. nov. and Paraprevotella xylaniphila sp. nov., members of the family ‘Prevotellaceae’ isolated from human faeces. Int J Syst Evol Microbiol 59, 1895–1900 (2009).

56. Wu, G.D. et al. Linking long-term dietary patterns with gut microbial enterotypes. Science 334, 105–108 (2011).

57. UniProt, C. UniProt: a hub for protein information. Nucleic Acids Res 43, D204–212 (2015).

58. Ye, J., McGinnis, S. & Madden, T.L. BLAST: improvements for better sequence analysis. Nucleic Acids Res 34, W6–9 (2006).

59. Fu, L., Niu, B., Zhu, Z., Wu, S. & Li, W. CD-HIT: accelerated for clustering the next-generation sequencing data. Bioinformatics 28, 3150–3152 (2012).

60. Hu, E.J., et al. Lora: Low-rank adaptation of large language models. arXiv preprint arXiv*:2106.09685* (2021).

61. Huang, P. et al. Gut microbial genomes with paired isolates from China illustrate probiotic and cardiometabolic effects. Cell Genom 4, 100559 (2024).

62. Carter, M.M. et al. Ultra-deep sequencing of Hadza hunter-gatherers recovers vanishing gut microbes. Cell 186, 3111–3124 e3113 (2023).

63. Zhu, J. et al. Over 50,000 Metagenomically Assembled Draft Genomes for the Human Oral Microbiome Reveal New Taxa. Genomics Proteomics Bioinformatics 20, 246–259 (2022).

64. Wibowo, M.C. et al. Reconstruction of ancient microbial genomes from the human gut. Nature 594, 234–239 (2021).

65. Liu, Y. et al. A genome and gene catalog of glacier microbiomes. Nat Biotechnol 40, 1341–1348 (2022).

66. Richardson, L. et al. MGnify: the microbiome sequence data analysis resource in 2023. Nucleic Acids Res 51, D753–D759 (2023).

67. Schubert, M., Lindgreen, S. & Orlando, L. AdapterRemoval v2: rapid adapter trimming, identification, and read merging. BMC Res Notes 9, 88 (2016).

68. Martin, M.J.E.j. Cutadapt removes adapter sequences from high-throughput sequencing reads. EMBnet. journal 17, 10–12 (2011).

69. Li, D., Liu, C.M., Luo, R., Sadakane, K. & Lam, T.W. MEGAHIT: an ultra-fast single-node solution for large and complex metagenomics assembly via succinct de Bruijn graph. Bioinformatics 31, 1674–1676 (2015).

70. Langmead, B. & Salzberg, S.L. Fast gapped-read alignment with Bowtie 2. Nat Methods 9, 357–359 (2012).

71. Li, H. et al. The Sequence Alignment/Map format and SAMtools. Bioinformatics 25, 2078–2079 (2009).

72. Kang, D.D. et al. MetaBAT 2: an adaptive binning algorithm for robust and efficient genome reconstruction from metagenome assemblies. PeerJ 7, e7359 (2019).

73. Chaumeil, P.A., Mussig, A.J., Hugenholtz, P. & Parks, D.H. GTDB-Tk: a toolkit to classify genomes with the Genome Taxonomy Database. Bioinformatics 36, 1925–1927 (2019).

74. Larralde, M. Pyrodigal: Python bindings and interface to Prodigal, an efficient method for gene prediction in prokaryotes. Journal of Open Source Software 7, 4296 (2022).

75. Steinegger, M. & Soding, J. MMseqs2 enables sensitive protein sequence searching for the analysis of massive data sets. Nat Biotechnol 35, 1026–1028 (2017).

76. Cantalapiedra, C.P., Hernández-Plaza, A., Letunic, I., Bork, P. & Huerta-Cepas, J. eggNOG-mapper v2: functional annotation, orthology assignments, and domain prediction at the metagenomic scale. Molecular biology and evolution 38, 5825–5829 (2021).

77. Feng, X. et al. Genomes of multicellular algal sisters to land plants illuminate signaling network evolution. Nat Genet 56, 1018–1031 (2024).

78. Mirdita, M. et al. ColabFold: making protein folding accessible to all. Nature methods 19, 679–682 (2022).

79. Cock, P.J. et al. Biopython: freely available Python tools for computational molecular biology and bioinformatics. Bioinformatics 25, 1422–1423 (2009).

80. Zhang, C., Shine, M., Pyle, A.M. & Zhang, Y. US-align: universal structure alignments of proteins, nucleic acids, and macromolecular complexes. Nat Methods 19, 1109–1115 (2022).

81. Tang, Q.Y., Ren, W., Wang, J. & Kaneko, K. The Statistical Trends of Protein Evolution: A Lesson from AlphaFold Database. Mol Biol Evol 39 (2022).

82. Katoh, K. & Standley, D.M. MAFFT multiple sequence alignment software version 7: improvements in performance and usability. Mol Biol Evol 30, 772–780 (2013).

83. Nguyen, L.T., Schmidt, H.A., von Haeseler, A. & Minh, B.Q. IQ-TREE: a fast and effective stochastic algorithm for estimating maximum-likelihood phylogenies. Mol Biol Evol 32, 268–274 (2015).

84. Letunic, I. & Bork, P. Interactive Tree Of Life (iTOL) v5: an online tool for phylogenetic tree display and annotation. Nucleic Acids Res 49, W293–W296 (2021).

85. Wiegand, I., Hilpert, K. & Hancock, R.E. Agar and broth dilution methods to determine the minimal inhibitory concentration (MIC) of antimicrobial substances. Nat Protoc 3, 163–175 (2008).

86. Jo, S., Kim, T., Iyer, V.G. & Im, W. CHARMM-GUI: a web-based graphical user interface for CHARMM. J Comput Chem 29, 1859–1865 (2008).

87. Case, D.A. et al. AmberTools. Journal of chemical information and modeling 63, 6183–6191 (2023).

88. Case, D.A., et al. AMBER 9. University of California, San Francisco 45 (2006).

89. Ryckaert, J.-P., Ciccotti, G. & Berendsen, H.J.C. Numerical integration of the cartesian equations of motion of a system with constraints: molecular dynamics of n-alkanes. Journal of computational physics 23, 327–341 (1977).

90. Wang, E. et al. Assessing the performance of the MM/PBSA and MM/GBSA methods. 10. Impacts of enhanced sampling and variable dielectric model on protein-protein Interactions. Phys Chem Chem Phys 21, 18958–18969 (2019).

