## Supplementary Information for "Unveiling the Evolution of Antimicrobial Peptides in Gut Microbes via Foundation Model-Powered Framework"

**Table of Contents**

**List of Abbreviations**

**Supplementary Fig. 1 |** Selection of model parameters for AMP discovery.

**Supplementary Fig. 2 |** Physicochemical properties of AMP sequences

**Supplementary Fig. 3 |** Violin diagrams of structural properties for AMP-containing proteins from the APD3 and LAMP2 databases (shown in orange and green, respectively) compared to those identified in this study (shown in pink).

**Supplementary Fig. 4 |** 500-ns MD simulation of AMP candidate (p1, p17 and variants) interacting with a Gram-positive bacterial membrane (G+PM).

**Supplementary Fig. 5 |** Molecular mechanics/generalized Born surface area (MM/GBSA) binding free energy calculations for AMP_c423 and AMP_c268 variants

**Supplementary Fig. 6 |** Detailed results of cytotoxicity of AMP candidates.

**Supplementary Fig. 7 |** Detailed results of hemolytic activity of AMP candidates selected based on clustering.

**Supplementary Fig. 8 |** SignalP results for p17.

**Supplementary Fig. 9 |** Orthogroups of AMP-containing proteins from various human gut metagenomics datasets.

**Supplementary Table 1 |** Comparison of classification performance on validation dataset: frozen vs. Fine-Tuned foundation models.

**Supplementary Table 2 |** Comparison of classification performance on validation dataset: end-to-end token classification vs. sequence and token classification

**Supplementary Table 3 |** Comparison of classification performance on validation dataset: linear classifier vs. LSTM classifier.

**Supplementary Table 4 |** Comparison of classification performance on validation dataset: combined loss vs. token loss only.

**Supplementary Table 5 |** Comparison of classification performance on validation dataset: CE loss vs. balanced CE loss.

**Supplementary Table 6 |** Comparison of classification performance on validation dataset: fine-tuning vs. LoRA.

**Supplementary Table 7** | Overview of published approaches for AMP Identification

**Supplementary Table 8 |** Performance of various approaches on identifying smORF AMPs.

**List of Abbreviations**

**AMP**	antimicrobial peptide

**smORF**	small open reading frame

**AMP-SEMiner**	Antimicrobial Peptide Structural Evolution Miner

**MSA**	multiple sequence alignment

**HMM**	hidden markov model

**GO**	gene ontology

**PPV**	predictive positive value

**PqqD**	coenzyme PQQ synthesis protein D

**LLM**	large langue model

**CE**	cross-entropy

**3D**	three-dimension

**MIC**	minimum inhibitory concentration

**CC50**	the 50% cytotoxic concentration

**HC50**	the 50% hemolytic concentration

**PI**	propidium iodide

**MD**	molecular dynamics

**G+PM**	gram-positive bacterial

**MM/GBSA**	Molecular mechanics/generalized Born surface area

**GRAMRA**	giant repository of AMP activities

**MAG**	metagenome-assembled genomes

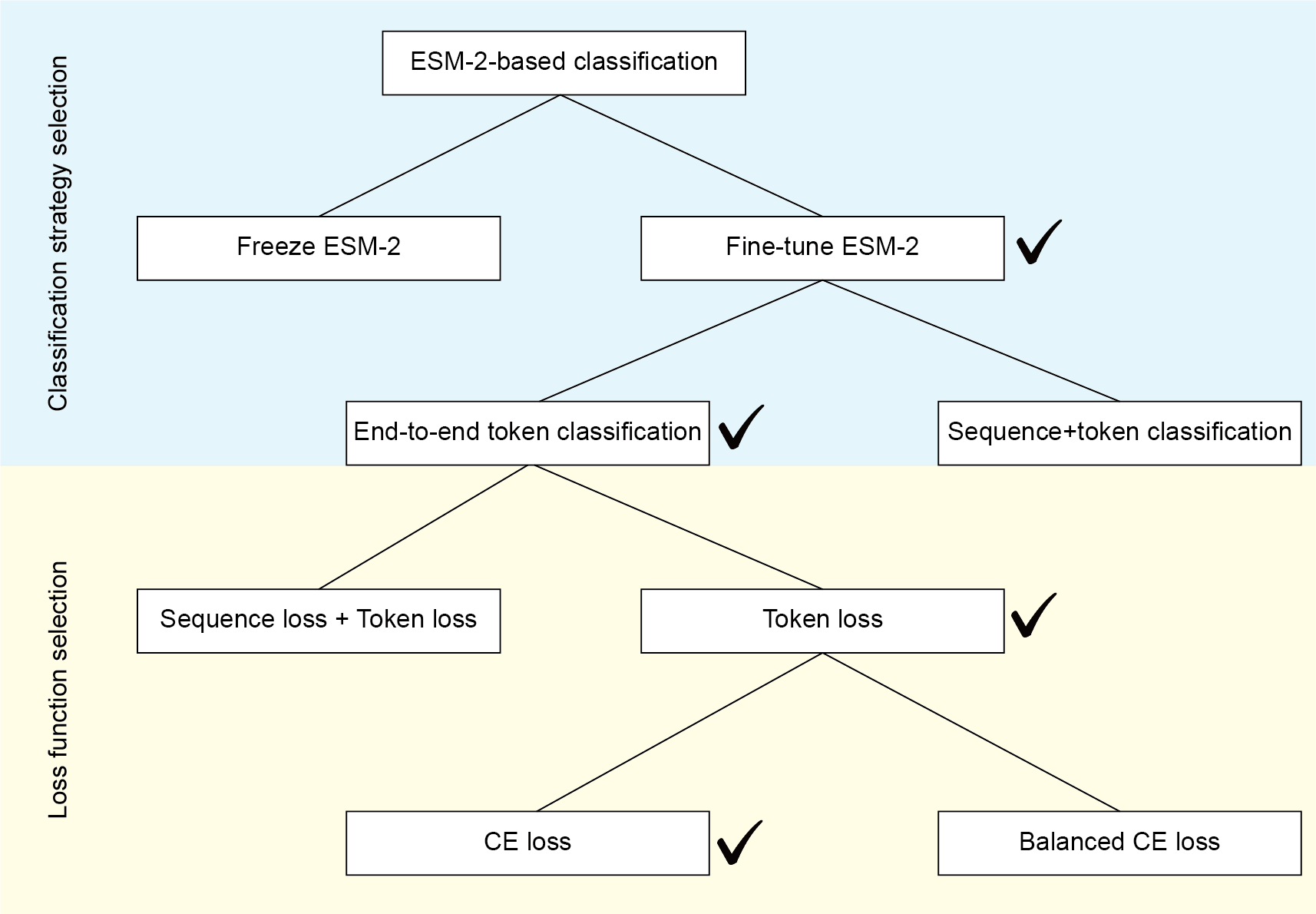

**Supplementary Fig. 1 |** Selection of model parameters for AMP discovery.

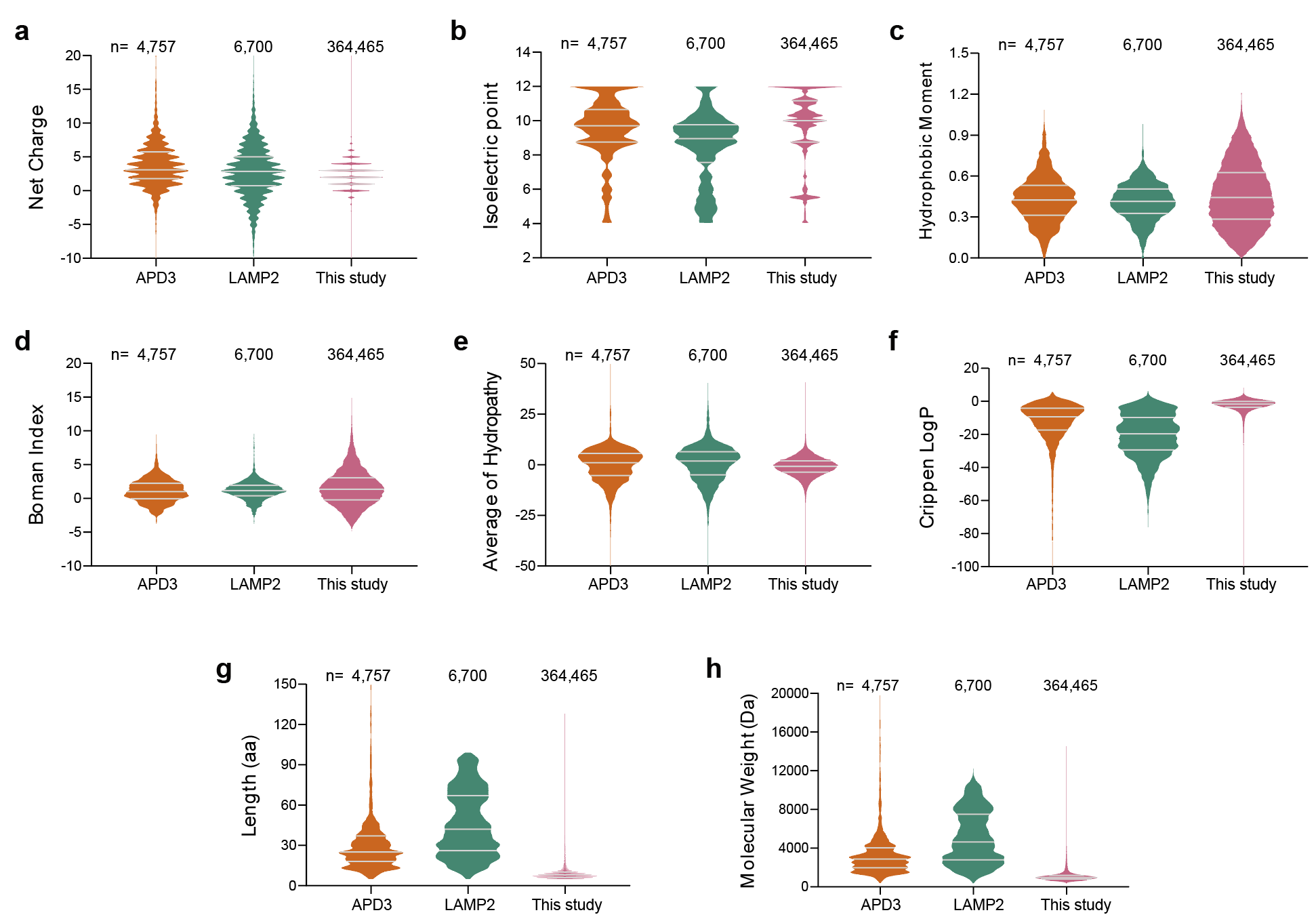

**Supplementary Fig. 2 |** Physicochemical properties of AMP sequences. a, Net charge; b, Isoelectric point; c, Hydrophobic moment; d, Boman index; e, Average hydropathy; f, Crippen LogP; g, Length (amino acids, aa); h, Molecular weight (Da).

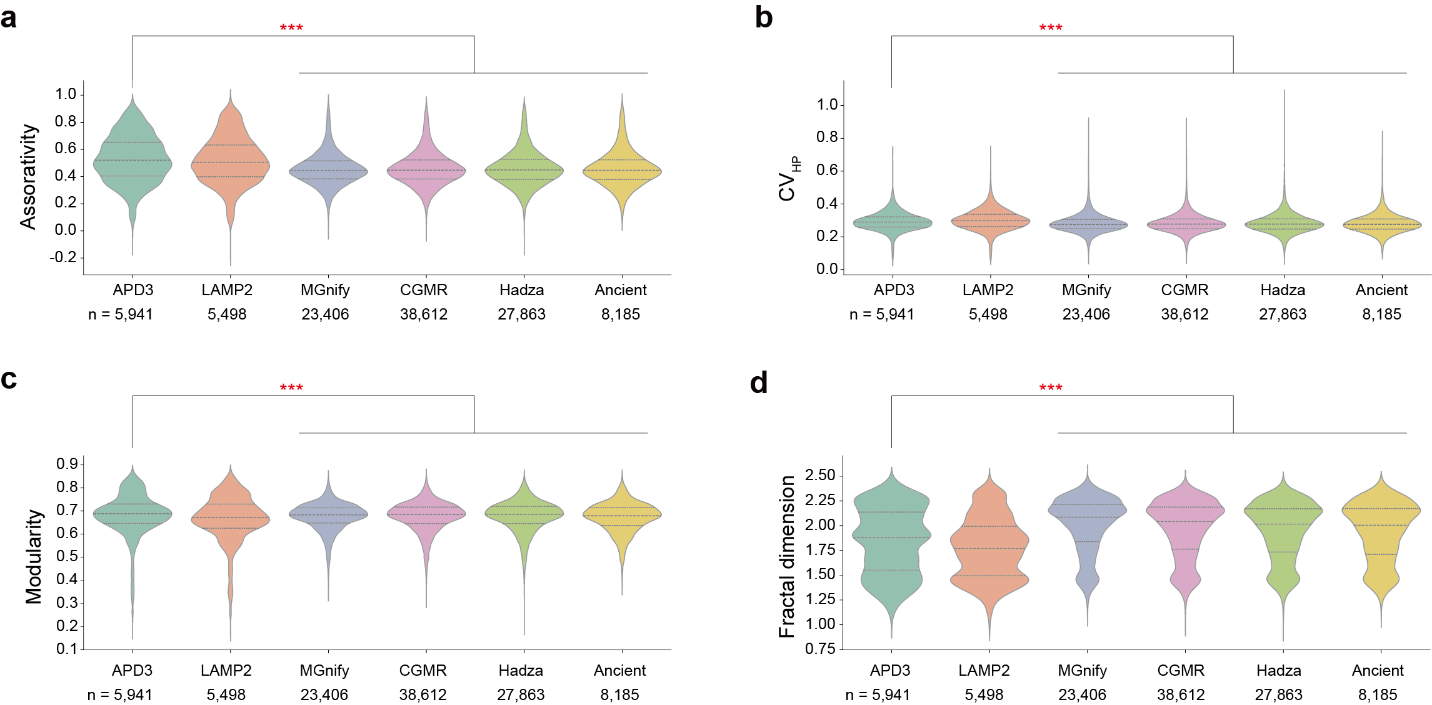

**Supplementary Fig. 3 |** Violin diagrams of structural properties for AMP-containing proteins from the APD3 and LAMP2 databases (shown in orange and green, respectively) compared to those identified in this study (shown in pink). a, Assortativity of the residue contact network, indicating the spatial segregation between ordered-and-disordered regions in protein sequences. b, CV_HP_, representing hydropathy variation to measure the hydrophilic-hydrophobic segregation in protein sequences, defined as the coefficient of variation of the filtered hydropathy profile of a sequence. c, Modularity of the residue contact network, showing the structural segmentation within protein networks. d, Fractal dimension, calculated using the box-counting method.

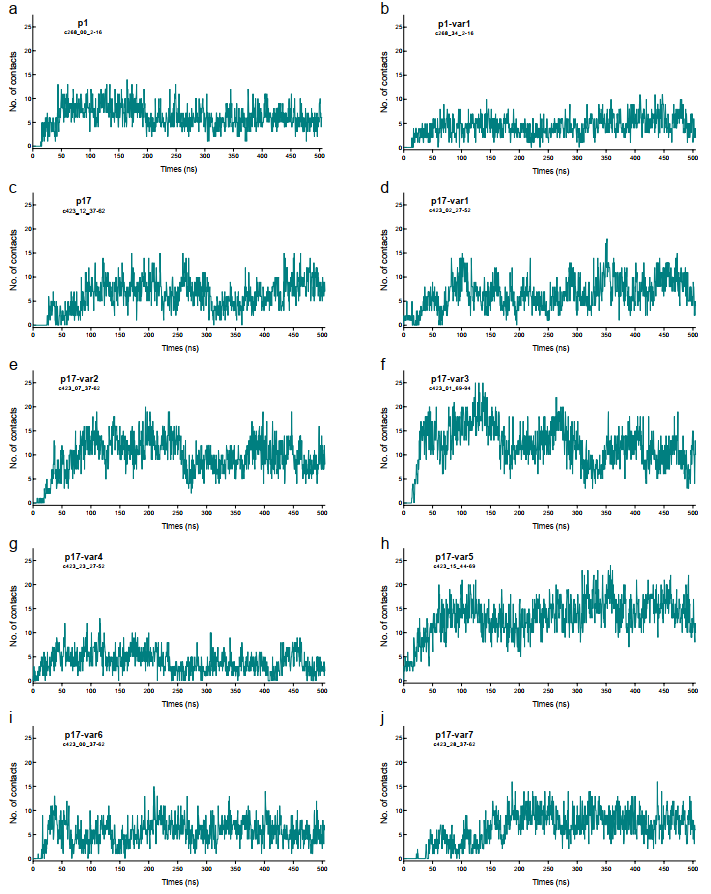

**Supplementary Fig. 4 |** A 500-ns MD simulation of p1(a), p1-var(b), p17(c), p17-var1(d), p17-var2(e), p17-var3(f), p17-var4(g), p17-var5(h), p17-var6(i) and p17-var7(j) variants interacting with a Gram-positive bacterial membrane (G+PM). A contact is defined as any heavy atom in the AMP being within 3Å of the membrane. The interaction resulted in stable peptide-membrane structures, demonstrating the antimicrobial mechanism as bacterial membrane disruption

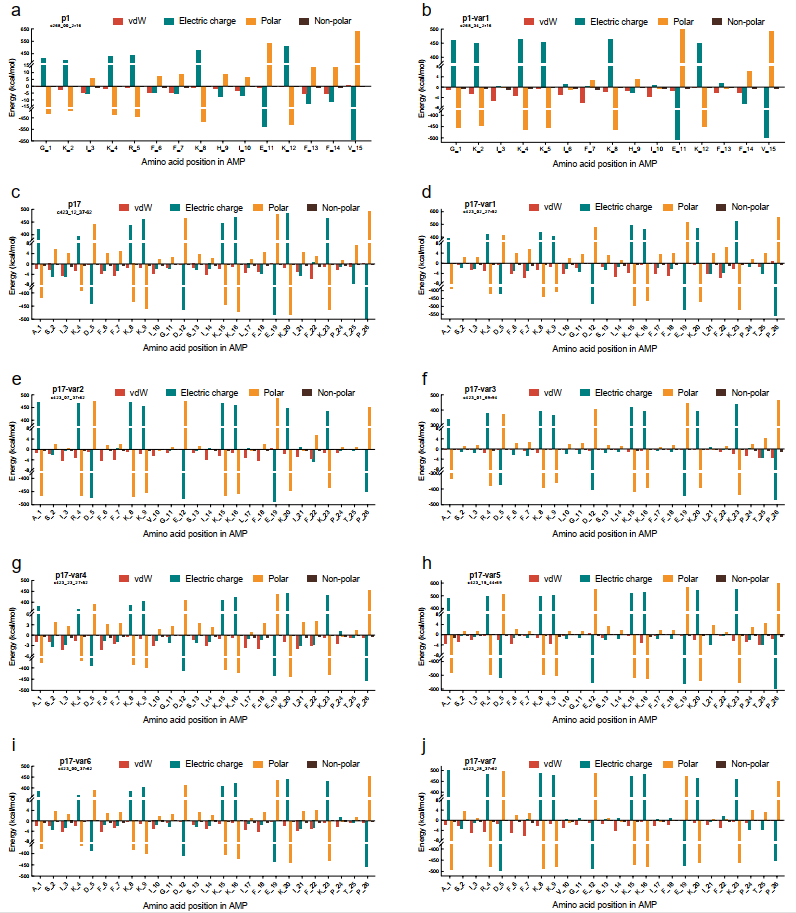

**Supplementary Fig. 5 |** Molecular mechanics/generalized Born surface area (MM/GBSA) binding free energy calculations for AMP_c423 and AMP_c268 variants

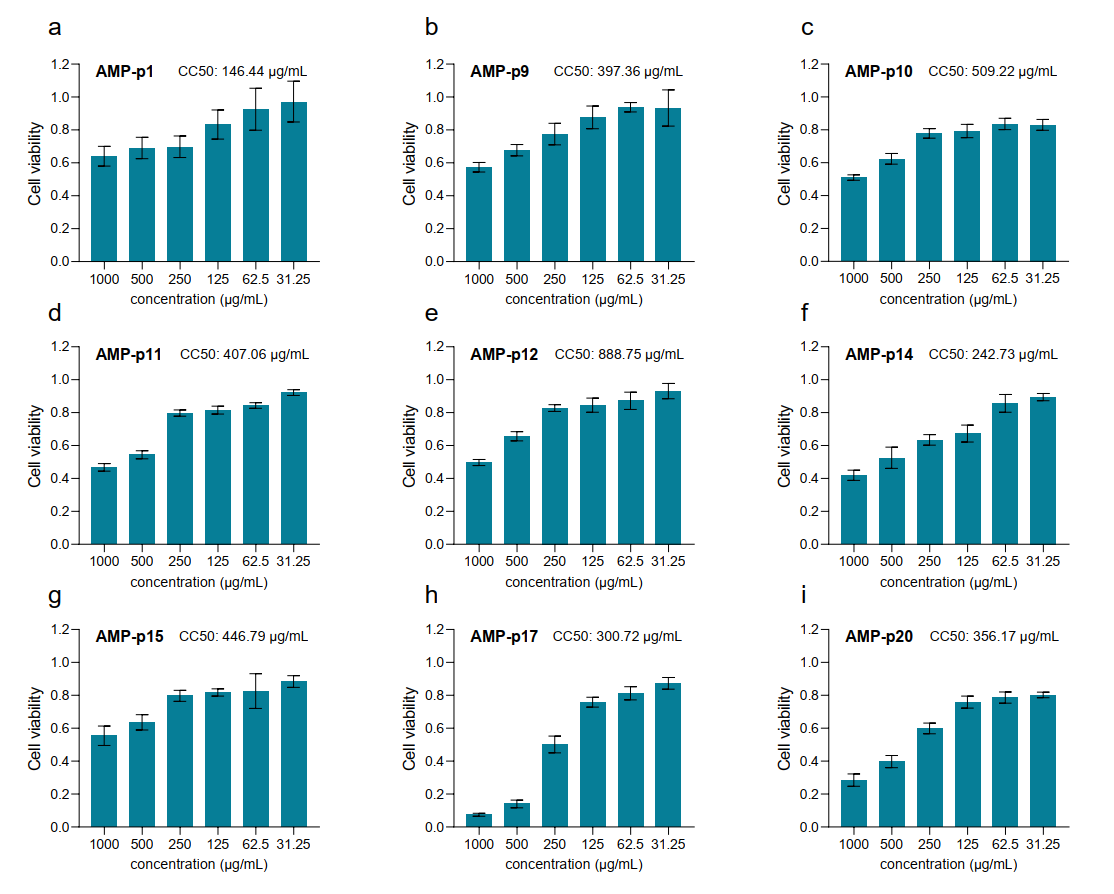

**Supplementary Fig. 6 |** Detailed results of cytotoxicity of AMP candidates.

a-r, Cytotoxicity results of peptides p1, p9, p10, p11, p12, p14, p15, p17 and p20 at different concentrations (1000 - 31.3 µg/mL). The CC50 values of the peptides are shown at the top of each panel.

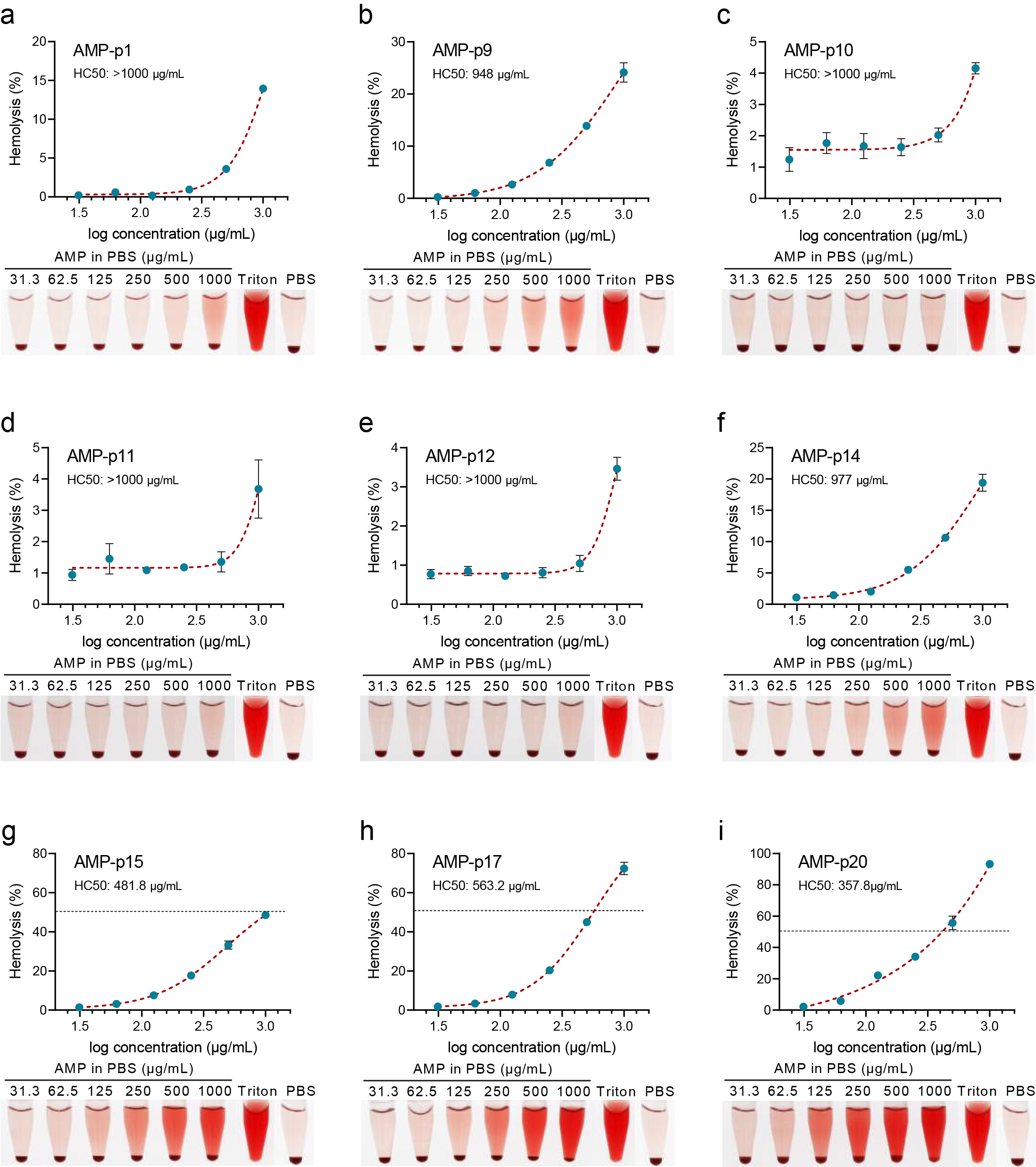

**Supplementary Fig. 7 |** Detailed results of hemolytic activity of AMP candidates selected based on clustering.

a-i, Hemolytic assay of peptides p1, p9, p10, p11, p12, p14, p15, p17, and p20, at different concentrations. The blue dots, from left to right, represent the concentrations 31.3, 62.5, 125, 250, 500, and 1000 µg/mL, which correspond to the concentrations of the tubes placed below. Bars are the average of n = 3 independent tests. Triton X-100 treated erythrocytes served as a positive control and the PBS treated erythrocytes acted as a negative control.

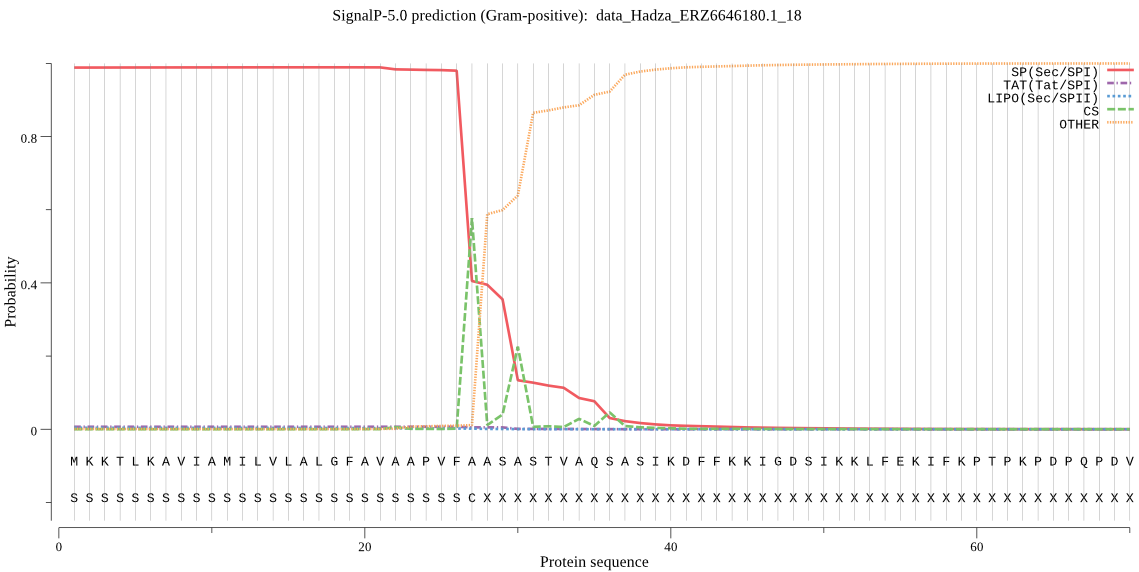
 **Supplementary Fig. 8 |** SignalP results for p17.

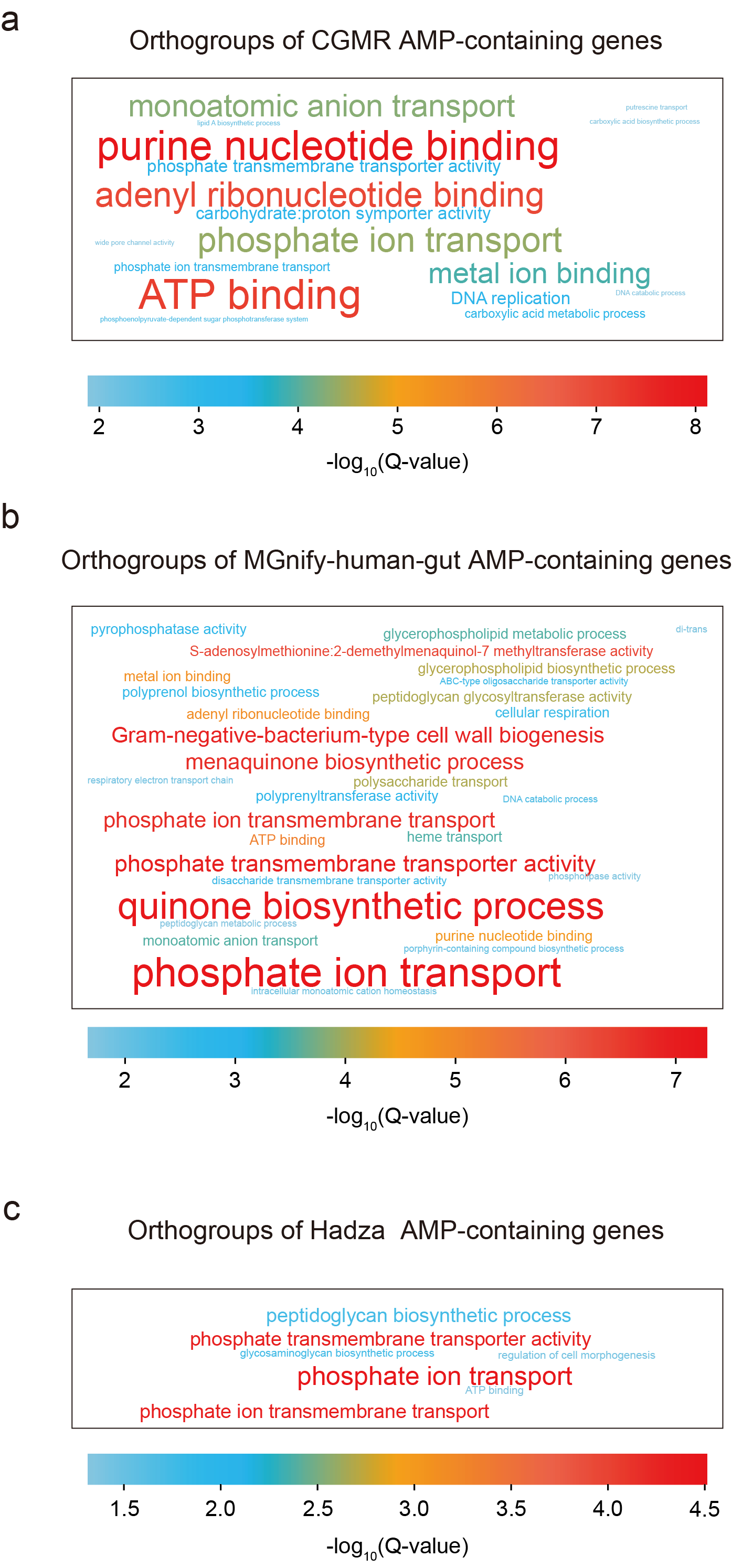

**Supplementary Fig. 9 |** Orthogroups of AMP-containing proteins from various human gut metagenomics datasets. a, CGMR dataset. b, MGnify-human-gut dataset. c, Hadza dataset.

**Supplementary Table 1 |** Comparison of classification performance on validation dataset: frozen vs. Fine-Tuned foundation models.

| **Classification level** | **Frozen or fine-tuned** | **Foundation model size** | **Training data size** | **Classification performance** | | | **Epoch** |
| --- | --- | --- | --- | --- | --- | --- | --- |
|  |  |  |  | **F1** | **Precision** | **Recall** |  |
| Sequence | Frozen | 150M | 20K | 0.473 | 0.765 | 0.342 | 22 |
| Sequence | Fine-tuned | 150M | 20K | **0.590** | 0.783 | **0.474** | 5 |
| Token | Frozen | 150M | 20K | 0.004 | **1.000** | 0.002 | 30 |
| Token | Fine-tuned | 150M | 20K | 0.534 | 0.833 | 0.395 | 12 |
| Sequence | Frozen | 650M | 2M | 0.690 | 0.840 | 0.586 | 15 |
| Sequence | Fine-tuned | 650M | 2M | 0.732 | 0.844 | 0.647 | 20 |
| Token | Frozen | 650M | 2M | 0.354 | 0.755 | 0.231 | 12 |
| Token | Fine-tuned | 650M | 2M | **0.823** | **0.887** | **0.768** | 12 |

**Supplementary Table 2 |** Comparison of classification performance on validation dataset: end-to-end token classification vs. sequence and token classification.

| **Classification strategy** | **Foundation model size** | **Training data size** | **Classification performance** | | | **Epoch** |
| --- | --- | --- | --- | --- | --- | --- |
|  |  |  | **F1** | **Precision** | **Recall** |  |
| Sequence+token classification | 650M | 2M | 0.674 | 0.746 | 0.615 | 20+50 |
| End-to-end token classification | 650M | 2M | **0.822** | **0.903** | **0.754** | 15 |

**Supplementary Table 3 |** Comparison of classification performance on validation dataset: linear classifier vs. LSTM classifier.

| **MLP** | **Foundation model size** | **Training data size** | **Classification performance** | | | **Epoch** |
| --- | --- | --- | --- | --- | --- | --- |
|  |  |  | **F1** | **Precision** | **Recall** |  |
| Linear classifier | 650M | 2M | **0.822** | **0.903** | **0.754** | 15 |
| LSTM classifier | 650M | 2M | 0.671 | 0.799 | 0.578 | 15 |

**Supplementary Table 4 |** Comparison of classification performance on validation dataset: combined loss vs. token loss only.

| **Loss function**  **strategy** | **Foundation model size** | **Training data size** | **Classification performance** | | | **Epoch** |
| --- | --- | --- | --- | --- | --- | --- |
|  |  |  | **F1** | **Precision** | **Recall** |  |
| Sequence loss + token loss | 650M | 2M | 0.571 | 0.753 | 0.459 | 13 |
| Token loss only | 650M | 2M | **0.819** | **0.903** | **0.750** | 13 |

**Supplementary Table 5 |** Comparison of classification performance on validation dataset: CE loss vs. balanced CE loss

| **Loss function**  **strategy** | **Foundation model size** | **Training data size** | **Classification performance** | | | **Epoch** |
| --- | --- | --- | --- | --- | --- | --- |
|  |  |  | **F1** | **Precision** | **Recall** |  |
| CE loss | 650M | 2M | **0.819** | **0.903** | **0.750** | 13 |
| Balanced CE loss | 650M | 2M | 0.494 | 0.431 | 0.578 | 13 |

**Supplementary Table 6 |** Comparison of classification performance on validation dataset: fine-tuning vs. LoRA

| **Fine-tuning strategy** | **Foundation model size** | **Training data size** | **Classification performance** | | | **Epoch** |
| --- | --- | --- | --- | --- | --- | --- |
|  |  |  | **F1** | **Precision** | **Recall** |  |
| Full fine-tune | 650M | 2M | 0.823 | **0.887** | 0.768 | 12 |
| LoRA  (output.dense, rank=8) | 650M | 2M | **0.836** | 0.883 | **0.794** | 12 |

**Supplementary Table 7** | Overview of published approaches for AMP Identification

| **No.** | **Approach** | **Journal** | **Year** | **Remark** |
| --- | --- | --- | --- | --- |
| 1 | SAMP^1^ | bioRxiv | 2024 | **Tested** |
| 2 | PGAT-ABPp^2^ | Bioinformatics | 2024 | **Tested** |
| 3 | deepAMPNet^3^ | PeerJ | 2024 | **Tested** |
| 4 | sAMPpred-GAT^4^ | Bioinformatics | 2023 | The required database (nrdb90) is unavailable |
| 5 | AMP-BERT^5^ | Protein Sci | 2023 | **Tested** |
| 6 | iAMPCN^6^ | Briefings in Bioinformatics | 2023 | **Tested** |
| 7 | panCleave^7^ | Cell Host & Microbe | 2023 | Only for human proteins. |
| 8 | Ma et at.^8^ | Nature Biotechnology | 2022 | **Tested** |
| 9 | AMPlify^9^ | BMC Genomics | 2022 | **Tested** |
| 10 | ampir^10^ | Bioinformatics | 2021 | **Tested** |
| 11 | amPEPpy 1.0^11^ | Bioinformatics | 2021 | **Tested** |
| 12 | Macrel^12^ | PeerJ | 2020 | **Tested** |
| 13 | AmpGram^13^ | International journal of molecular sciences | 2020 | **Tested** |
| 14 | Deep-AMPEP30^14^ | Molecular Therapy - Nucleic Acids | 2020 | Only for proteins with length <= 30 aa. |
| 15 | IAMPE^15^ | Journal of Chemical Information and Modeling | 2020 | Code unavailable. |
| 16 | AMPfun^16^ | Briefings in Bioinformatics | 2020 | Not supported for large input data sets. |
| 17 | APIN^17^ | BMC Genomics | 2019 | No trained model provided. |
| 18 | AMAP^18^ | Computers in Biology and Medicine | 2019 | Code unavailable. |
| 19 | AMP Scanner v2^19^ | Bioinformatics | 2018 | Not supported for large input data sets. |
| 20 | AmPEP^20^ | Scientific Reports | 2018 | Not supported for large input data sets. |
| 21 | iAMPpred^21^ | Scientific Reports | 2017 | Code unavailable. |

**Supplementary Table 8 |** Performance of various approaches on identifying smORF AMPs.

| **Approach** | **APD3 smORFs (254, 4.3%)** | **LAMP2 smORFs (942,17.2%)** |
| --- | --- | --- |
|  | **Recall** | **Recall** |
| Macrel^24^ | 0.689 | 0.207 |
| AmpGram^45^ | 0.917 | 0.832 |
| ampir^46^ | 0.177 | 0.587 |
| amPEPpy 1.0^47^ | 0.764 | 0.387 |
| Ma et al.^25^ | 0.819 | 0.737 |
| AMPlify-balanced^48^ | 0.890 | 0.381 |
| AMPlify-imbalanced^49^ | 0.878 | 0.323 |
| iAMPCN^50^ | 0.913 | 0.732 |
| AMP-BERT^26^ | 0.925 | 0.901 |
| PGAT-ABPp^28^ | 0.378 | 0.238 |
| deepAMPNet^27^ | **0.976** | **0.973** |
| SAMP^51^ | 0.752 | 0.554 |
| Our approach | 0.744 | 0.238 |
|  | 0.657 | 0.209 |
|  | 0.685 | 0.229 |

References

1.	Feng, J. *et al.* SAMP: Identifying Antimicrobial Peptides by an Ensemble Learning Model Based on Proportionalized Split Amino Acid Composition. Preprint at https://doi.org/10.1101/2024.04.25.590553 (2024).

2.	Hao, Y., Liu, X., Fu, H., Shao, X. & Cai, W. PGAT-ABPp: harnessing protein language models and graph attention networks for antibacterial peptide identification with remarkable accuracy. *Bioinformatics* **40**, btae497 (2024).

3.	Zhao, F. *et al.* deepAMPNet: a novel antimicrobial peptide predictor employing AlphaFold2 predicted structures and a bi-directional long short-term memory protein language model. *PeerJ* **12**, e17729 (2024).

4.	Yan, K., Lv, H., Guo, Y., Peng, W. & Liu, B. sAMPpred-GAT: prediction of antimicrobial peptide by graph attention network and predicted peptide structure. *Bioinformatics* **39**, btac715 (2023).

5.	Lee, H., Lee, S., Lee, I. & Nam, H. AMP‐BERT: Prediction of antimicrobial peptide function based on a BERT model. *Protein Science* **32**, e4529 (2023).

6.	Xu, J. *et al.* iAMPCN: a deep-learning approach for identifying antimicrobial peptides and their functional activities. *Briefings in Bioinformatics* **24**, bbad240 (2023).

7.	Maasch, J. R. M. A., Torres, M. D. T., Melo, M. C. R. & De La Fuente-Nunez, C. Molecular de-extinction of ancient antimicrobial peptides enabled by machine learning. *Cell Host & Microbe* **31**, 1260-1274.e6 (2023).

8.	Ma, Y. *et al.* Identification of antimicrobial peptides from the human gut microbiome using deep learning. *Nat Biotechnol* **40**, 921–931 (2022).

9.	Li, C. *et al.* AMPlify: attentive deep learning model for discovery of novel antimicrobial peptides effective against WHO priority pathogens. *BMC Genomics* **23**, 77 (2022).

10.	Fingerhut, L. C. H. W., Miller, D. J., Strugnell, J. M., Daly, N. L. & Cooke, I. R. ampir: an R package for fast genome-wide prediction of antimicrobial peptides. *Bioinformatics* **36**, 5262–5263 (2021).

11.	Lawrence, T. J. *et al.* amPEPpy 1.0: a portable and accurate antimicrobial peptide prediction tool. *Bioinformatics* **37**, 2058–2060 (2021).

12.	Santos-Júnior, C. D., Pan, S., Zhao, X.-M. & Coelho, L. P. Macrel: antimicrobial peptide screening in genomes and metagenomes. *PeerJ* **8**, e10555 (2020).

13.	Burdukiewicz, M. *et al.* Proteomic Screening for Prediction and Design of Antimicrobial Peptides with AmpGram. *IJMS* **21**, 4310 (2020).

14.	Yan, J. *et al.* Deep-AmPEP30: Improve Short Antimicrobial Peptides Prediction with Deep Learning. *Molecular Therapy - Nucleic Acids* **20**, 882–894 (2020).

15.	Kavousi, K. *et al.* IAMPE: NMR-Assisted Computational Prediction of Antimicrobial Peptides. *J. Chem. Inf. Model.* **60**, 4691–4701 (2020).

16.	Chung, C.-R., Kuo, T.-R., Wu, L.-C., Lee, T.-Y. & Horng, J.-T. Characterization and identification of antimicrobial peptides with different functional activities. *Briefings in Bioinformatics* **21**, 1098–1114 (2020).

17.	Su, X., Xu, J., Yin, Y., Quan, X. & Zhang, H. Antimicrobial peptide identification using multi-scale convolutional network. *BMC Bioinformatics* **20**, 730 (2019).

18.	Gull, S., Shamim, N. & Minhas, F. AMAP: Hierarchical multi-label prediction of biologically active and antimicrobial peptides. *Computers in Biology and Medicine* **107**, 172–181 (2019).

19.	Veltri, D., Kamath, U. & Shehu, A. Deep learning improves antimicrobial peptide recognition. *Bioinformatics* **34**, 2740–2747 (2018).

20.	Bhadra, P., Yan, J., Li, J., Fong, S. & Siu, S. W. I. AmPEP: Sequence-based prediction of antimicrobial peptides using distribution patterns of amino acid properties and random forest. *Sci Rep* **8**, 1697 (2018).

21.	Meher, P. K., Sahu, T. K., Saini, V. & Rao, A. R. Predicting antimicrobial peptides with improved accuracy by incorporating the compositional, physico-chemical and structural features into Chou’s general PseAAC. *Sci Rep* **7**, 42362 (2017).
